# Diversification of the type IV filament super-family into machines for adhesion, secretion, DNA transformation and motility

**DOI:** 10.1101/576694

**Authors:** Rémi Denise, Sophie S Abby, Eduardo PC Rocha

## Abstract

Processes of molecular innovation require tinkering and co-option of existing genes. How this occurs in terms of molecular evolution at long evolutionary scales remains poorly understood. Here, we analyse the natural history of a vast group of membrane-associated molecular systems in Bacteria and Archaea – type IV filament super-family (TFF-SF) – that diversified in systems involved in flagellar or twitching motility, adhesion, protein secretion, and DNA natural transformation. We identified such systems in all phyla of the two domains of life, and their phylogeny suggests that they may have been present in the last universal common ancestor. From there, two lineages, a Bacterial and an Archaeal, diversified by multiple gene duplications of the ATPases, gene fission of the integral membrane platform, and accretion of novel components. Surprisingly, we find that the Tad systems originated from the inter-kingdom transfer from Archaea to Bacteria of a system resembling the Epd pilus. The phylogeny and content of ancestral systems suggest that initial bacterial pili were engaged in cell motility and/or DNA transformation. In contrast, specialized protein secretion systems arose much later, and several independent times, in natural history. All these processes of functional diversification were accompanied by genetic rearrangements with implications for genetic regulation and horizontal gene transfer: systems encoded in fewer loci were more frequently exchanged between taxa. Overall, the evolutionary history of the TFF-SF by itself provides an impressive catalogue of the variety of molecular mechanisms involved in the origins of novel functions by tinkering and co-option of cellular machineries.

## Introduction

New complex forms, functions, and molecular systems arise by co-option of elements that may have evolved to tackle different adaptive needs [1]. At the molecular level, this involves tinkering pre-existing molecular structures by diverse processes including mutation, recombination, gene fusion and fission [2]. These variants are ultimately subject to natural selection and may eventually become fixed in populations [3]. In Prokaryotes, this is facilitated by the constant income of novel genetic information by horizontal gene transfer [4–6]. Complex adaptations can evolve through series of small adaptive steps. For example, metabolic networks evolve stepwise to accommodate novel reactions at their edges [7]. Innovation may also arise by processes of neo-functionalization or sub-functionalization following the duplication of genes encoding proteins with multiple functions or the acquisition of a genetic system with homologs in the genome. This mechanism may have been at the origin of the bacterial flagellum [8]. It has been proposed that non-adaptive processes acting on redundant genes in species with small effective population sizes provide substrates for the secondary evolution of complex traits by natural selection [9]. Less is known about how key macromolecular complexes can evolve novel functions, or specialize in one of several initial functions, thanks to a combination of mutational processes and horizontal gene transfer [10].

The appendages of bacteria are striking examples of functional diversification. They are complex macromolecular machineries encoded by many genes and spanning several cellular compartments that can evolve towards novel functions. For example, the type III protein secretion system (T3SS) evolved from the secretion apparatus of the bacterial flagellum [11], the T4SS from the conjugation apparatus [12], and the T6SS possibly from co-option of phage structures [13, 14]. A particularly remarkable illustration of these processes is provided by the type IV filament super-family (TFF-SF) of bacterial and archaeal systems that include the type II protein secretion system (T2SS), the type IVa pilus (T4aP), the type IVb pilus (T4bP), the mannose-sensitive hemagglutinin pilus (MSH), the tight adherence pilus (Tad), the competence pilus (Com), and the type IV-related pili in Archaea (Archaeal-T4P), that includes the archaeal flagella (Archaellum). These systems have core homologous components, sometimes in multiple copies, and present similarities in terms of macro-molecular architecture (Fig. 1) [15–17]. They include AAA+ ATPases, among which the T4aP PilT is the most powerful molecular motor known [18], an integral (cytoplasmic) membrane (IM) platform, and a prepilin peptidase that matures a set of specific pilins or pseudo-pilins (in T2SS) [19]. Bacteria with two cell membranes (diderms) also encode a secretin that forms an outer membrane pore [20]. Other proteins of these systems are either specific for each system or evolve too fast to allow the inference of homology among all variants.

**Fig 1.**
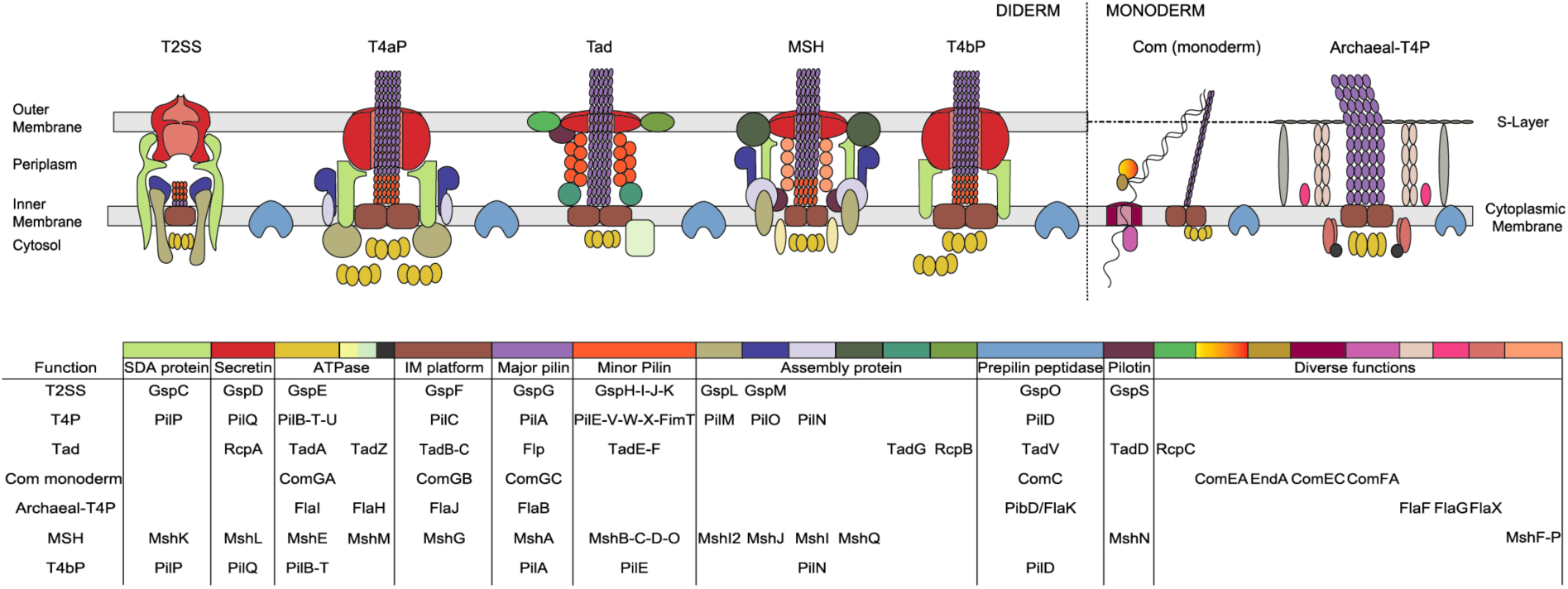
Schematic representation of the different systems and associated genes. Homologous components are represented in the same colour. The table below the drawing indicates the colour code and the name of the different components in each type of system. SDA stand for secretin-dynamic associated and IM stand for integral membrane. For the Archaeal-T4P the representation of the systems is based on the representation of the Archaellum and the genes mentioned in the legend are the names of the genes used in the literature not the arCOG names.

The TFF-SF nanomachines assemble filaments composed of subunits with an N-terminal sequence motif named class III signal peptide, generically named type IV pilins [17]. These systems are involved in functions typically associated with extracellular pili in Prokaryotes, including adherence, cell-cell attachment, the formation of biofilms, and are exploited by phages for cell infection [21–23]. The T4aP, T2SS, T4bP, and Tad, are also important virulence factors in pathogenic bacteria [24–29]. Nevertheless, and in spite of their homology, the different families of the TFF-SF have evolved specific biological functions. The T4aP and T4bP allow bacteria to move by twitching motility (a form of surface movement promoted by repeated cycles of extension-retraction of the pilus) [30, 31]. Tad are involved in efficient adherence to abiotic surfaces facilitating the formation of biofilms [32]. T2SS secrete proteins from the periplasm across the outer membrane [19]. Some Com, T4aP, and Archaeal-T4P facilitate the uptake of extracellular DNA into the cell [33, 34]. In Bacteria, these systems are by far the most frequent appendages involved in natural transformation. Archaeal-T4Ps include the archaellum involved in motility by rotation of the appendage, pili involved in sugar uptake (Bindosome, or Bas), the *Ups* pilus involved in establishing cell-cell contacts to enable DNA repair under stress conditions, and several pili with poorly characterized functions [35–37].

The functional diversification of the super-family is not clade-specific since different types of systems are present in the same clades. This suggests frequent horizontal transfer and/or an ancient origin of the super-family. T4aP and the Tad pilus can be found in most bacterial phyla [38, 39] and Archaeal-T4Ps in most Archaea [37]. The T2SS, T4bP and MSH have only been described in diderms [40, 41]. The distribution of the competence pilus is poorly known. Within diderms, a T4aP is often necessary but not sufficient for competence, which requires a specific machinery for DNA transport across the inner membrane into the cytoplasm [34]. Furthermore, a single T4aP can play other roles in addition to natural transformation (e.g. adhesion, motility, aggregation). In monoderms (bacteria with a single cell membrane), it is also currently unknown how many Competence pilus (ComM) systems are involved in this process. In summary, the TFF-SF has diversified into several different functions by co-option processes using a common set of homologous components identifiable across Prokaryotes.

Previous studies dedicated to the evolution of the AAA+ ATPases, Tad, T4aP and the T2SS date from the previous decade [15, 39, 42, 43], when data were scarce and phylogenetic methods less sophisticated. Archaeal systems were studied in detail recently [37, 44], but independently of the evolution of bacterial systems. More recent works only briefly studied the phylogenies of some of the components of these systems [45]. Importantly, there is a lack of studies integrating all the systems and all available genomic data, a pre-requisite to understand the processes of functional diversification of the super-family. Here, we identified the typical systems of TFF-SF and their variants using specific annotation tools on all complete genomes of Prokaryotes. These systems were analysed using phylogenetic techniques to characterize the history of the TFF-SF, clarify the relationships among its members, and decipher the molecular evolution mechanisms underlying its functional diversification. Finally, we characterized their genetic organizations, and how they relate to the rates of horizontal gene transfer. This integrative analysis provided a consistent scenario for the diversification of the super-family involving processes of duplication, fission, transfer, accretion, and mutation.

## Results

### Relations of homology between the key components of the machineries

We started our study by building stringent MacSyFinder models [46] for the identification of the systems of the TFF-SF in the complete genomes of Bacteria and Archaea. Briefly, these models give a detailed account of the genetic composition and organization of the systems and are used by MacSyFinder to identify novel occurrences of the systems in genomes using HMM protein profiles and a list of rules concerning genetic organization. At this stage, our goal was to identify systems with high stringency that would allow us to perform phylogenetic and comparative genomics analyses. We adapted previously published models of T4aP (including the Com systems of diderms), T2SS, and Tad [41, 47], to which we incorporated additional components and stricter rules in terms of genetic composition and organization (Fig S1). We used the available literature to produce equivalent models, and associated hidden Markov models (HMM) protein profiles, for the Com pilus of monoderms designated as ComM, and for the Archaeal-T4P. For the latter, we used 66 arCOGs identified from [37], after a step of re-analysis of the initial 191 arCOGs to remove redundancy. We could not build equivalent models for T4bP and MSH systems at this point because too few systems were described in the literature. This resulted in five initial models, of which two are novel, including 154 HMM protein profiles, of which 17 are novel (Table S1).

To establish precisely the relations of homology between the components of the different systems, we made pairwise profile-profile alignments of their HMM protein profiles using HHsearch v3.0.3 [48] (p-value threshold of 0.001). These alignments are very sensitive and highlight more distant relations of homology than typical sequence alignment methods [48]. We obtained a graph with 10 components (sets of connected nodes), representing the significant relations of reciprocal similarity between the profiles (Fig. 2). The five largest components include the proteins known to be homologous and represent each individual key function: secretins, prepilin peptidases, ATPases, integral membrane platforms (IM platform), pilins (major and minor). The ATPase component includes TadZ, a protein from another subfamily of P-loop ATPases (FleN) with an atypical Walker_-_A motif that retains ATP binding capacity while displaying low ATPase activity [49, 50]. It localizes at the pole at early stages of pili biogenesis and functions as a hub for recruiting other Tad pili components, contrary to the ATPases involved in pilus assembly or retraction.

**Fig 2.**
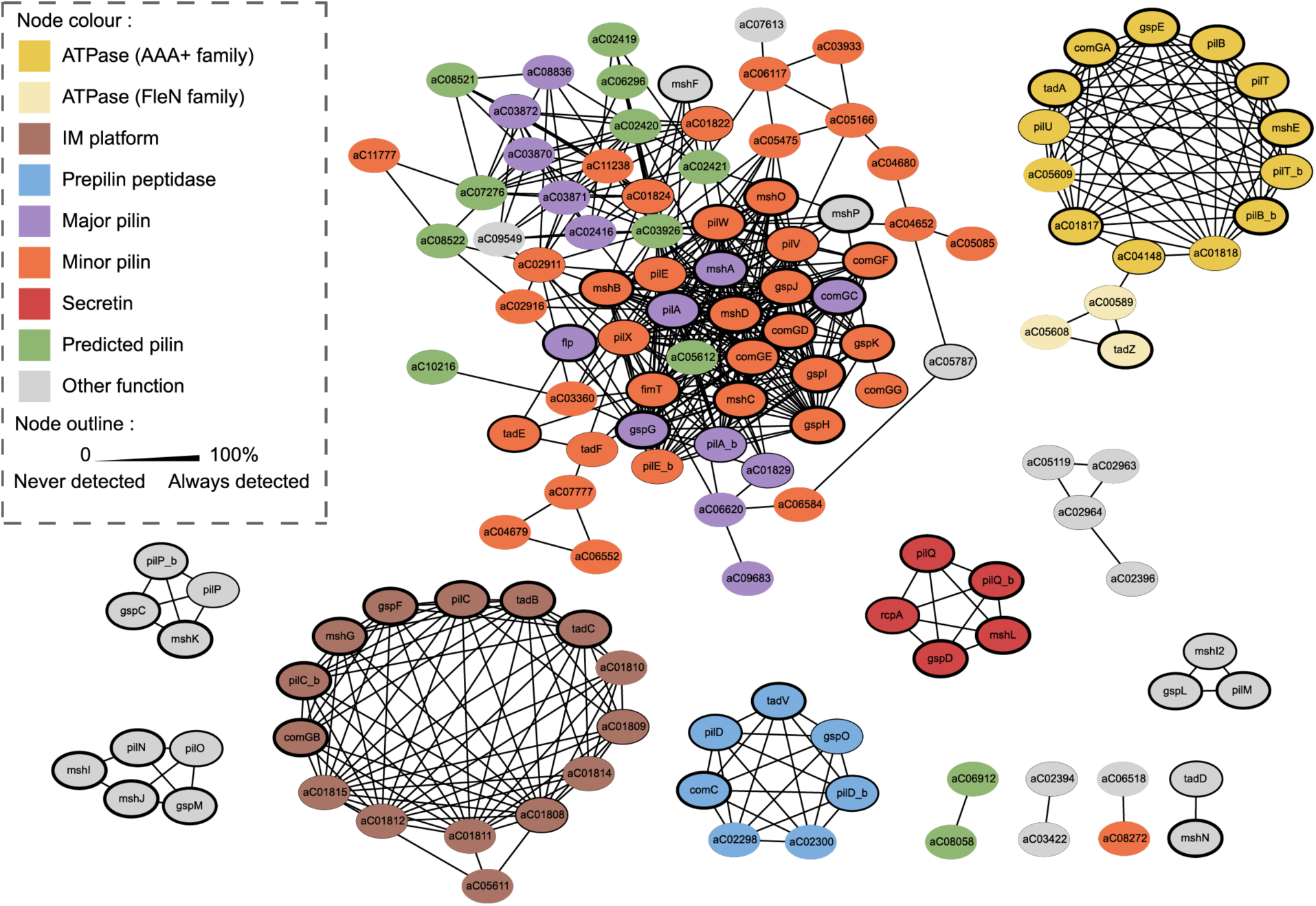
Results of the HMM-HMM alignments (HHSearch) between all the components of the TFF-SF. The colour of the nodes represents the known or predicted function of the protein. The outline of the node is proportional to the presence of the profile in the detected systems: the thinner the outline is, the less frequent the profile is found in the systems. The size of the outlines is proportional to the frequency of the profiles in the detected systems (thicker outlines indicate higher frequencies).

These results establish a precise and extensive network of sequence similarity between the key components of the systems of the TFF-SF, systematizing previous descriptions. The largest component of the graph includes the major and minor pilins. These proteins are small and very diverse. Many HMM profiles have been built for their correct identification, especially in Archaea [37]. Their profile-profile alignments suggest that these pilins are all evolutionarily related. It is for the moment unclear if all these archaeal pilin protein profiles are associated with families with different functions, but they are often similar and their profiles match the same proteins. This suggests that future work could reduce their number and produce a more parsimonious set of profiles for annotation, eventually revealing broader functional groups.

### The phylogenies of the components of the type IV filament super-family

The presence of homologs of the major functional components of the TFF-SF in most types of systems raises the question of how their functional diversification took place from a common ancestor. To study this, we added to the models described above a very simple generic model to identify all systems that have a minimal number of essential components (the ATPase, the integral membrane platform, and a major pilin) (Fig. S1). The search for systems using the MacSyFinder models resulted in the identification of 6652 systems in 3700 genomes (1486 species) (Fig S2), of which 1584 were classed as generic systems, reflecting the conservative character of the initial models. This dataset was too large to analyse using sophisticated phylogenetic methods and included many systems that were very similar, e.g. from different strains of the same species. We reduced this redundancy by clustering very similar systems. We then picked one representative per cluster, thus preserving most of the diversity of the dataset. In this process, we prioritized the inclusion of experimentally validated systems, including MSH (1), and T4bP (5) for which models were not available (see Methods). This non-redundant set contains 309 representative systems (33 T4aP, 47 Archaeal-T4P, 29 T2SS, 5 T4bP, 1 MSH, 31 ComM, 72 Tad, 101 generic) (Table S2). Hence, the systems used in the subsequent analyses are associated with a (sometimes large) number of other very similar systems that are from the same cluster.

We inferred the phylogeny of each of the five key protein components (AAA+ ATPase, IM platform, major pilin, secretin and prepilin peptidase) by maximum likelihood with IQ-Tree [51]. We made ten reconstructions per component with the most thorough mode of topological search to account for the stochasticity of the method. The detailed analysis of key events revealed by these trees can be found in Table S3 (the trees themselves are in Table S4). The ATPase trees are very well supported at most of the key nodes, they are consistent across replicated inferences, and they clearly separate the different types of systems (Fig. S3). The trees include two system-specific duplication events of the ATPases, one ancestral to the large clade including T4aP, T4bP, MSH, T2SS, and ComM (PilT/PilB), and another within a clade of T4aP (PilT/PilU). The IM platform also discriminates the different systems. It includes the well-known paralogs TadB/TadC in the Tad system and some duplications in the Archaea. Apart from these duplications that span large numbers of systems, there were other duplicates in the systems that were rare (present in 8% of the representatives’ dataset) and dealt with a de-replication procedure (see Methods). The prepilin peptidase tree is poorly supported and shows scattered distribution of the different types of systems. Since prepilin peptidases from one type of systems can be used by different systems [52–54], we have excluded them from further analyses. The secretin and major pilin trees have some poorly supported branches, but they separate well the different systems. Overall, the protein components’ trees show that the ATPase, the IM platform and the major pilin are phylogenetic markers that provide significant information about the evolution of the super-family. The secretin tree, even if relatively well supported, is less informative to infer the global evolutionary scenario because this component is absent from several types of systems (those from bacterial monoderms and archaea).

### The root of the type IV filament super-family

The ATPase tree is the only one that can be rooted, since this is the only ubiquitous component with well-conserved homologs in distinct machineries [42, 43]. We selected the family of FtsK proteins as an outgroup to root the tree. It represents an ideal candidate outgroup because it is very conserved in sequence, single-copy, and present in most bacterial phyla. It has a close homolog, HerA, which is the sister-clade among the family of P-loop ATPases from which it diverged concomitantly with the archaeal–bacterial division after the last universal common ancestor [43]. Furthermore, FtsK is essential (for chromosome segregation [55, 56]) and shows little evidence of horizontal transfer [43]. We retrieved the sequences of FtsK from a previous study [12], aligned them with the ATPase sequences of the investigated systems, and inferred a maximum likelihood tree. This tree shows that the FtsK sequences are monophyletic (100% UF-Boot support) and branch between two large clades: the Tad and Archaeal-T4P on one side (100% UF-boot) and a clade grouping the T2SS, T4aP, ComM and T4bP on the other side (100% UF-boot) (Fig S4). The overall rooted topology is very similar to that of the unrooted tree in eight out of ten trees (Table S3). The inclusion of the ubiquitous ATPase of T4SS (VirB4) as an outgroup with FtsK also showed a split between the archaeal and the bacterial branches of the tree (Fig S5). This confirms that this ATPase family is also an outgroup of the TFF-SF. We rooted the trees of the IM platform and major pilin using the root of the ATPase trees, since all three proteins showed a consistent split between Tad/T4P-Archaea on one side and the remaining systems on the other (Figs S6, S7).

The analysis of gene duplications provides additional information on the possible roots of the super-family phylogenetic tree because placing duplication events on a tree corresponds to set as ancestral the node where the duplications occurred, and as descendants those with the duplication [57, 58]. The duplications of the ATPases exclude the root from the group T4aP, T4bP, ComM, MSH and T2SS. The duplication of the IM platform in the Tad system, also present in some Archaeal-T4P, excludes the root from within these groups. Hence, the analyses of duplication events are consistent with the root as defined above by the tree of ATPases.

### Producing a concatenate tree

Since the two major components (ATPase and IM platform) have phylogenetic trees that are broadly consistent (Table S3), we computed a phylogenetic tree of their concatenate using a partition model (best model for each gene partition, as computed by IQ-Tree). The major pilin was excluded from the concatenate because it shows less consistent and less supported topologies. Concatenation required the use of a procedure to deal with paralogs (to have one marker per component per system). For paralogs present in a few taxa, we chose in each system the protein most similar in sequence to the most closely related systems lacking paralogs (see Methods). For the ATPases, we used PilB, because this ATPase is responsible for the assembly of the pilus, which is a function that is essential in all families, contrary to the function of PilT/PilU (retraction). There was no good argument to pick TadB or TadC platform proteins and we therefore made phylogenetic reconstructions with each of them in parallel (Fig. 3, S10). As expected, the concatenate trees showed relationships similar to those of the ATPase and the IM platform. We then tested the congruence between the two concatenate trees (PilB/TadB and PilB/TadC) and the individual protein alignments (PilB and TadB for the first, and PilB and TadC for the second) using the AU test implemented in IQ-Tree (v1.6.7.2) [59]. This showed that the best trees of each individual protein were not significantly different from the best tree of the concatenate (p>0.05, Table S7). Furthermore, after correction for multiple comparisons, only two of the 40 comparisons between the individual trees and the concatenate tree were significantly incongruent. Overall, these analyses strongly suggest that the TFF-SF derived from an ancestral system, which diversified initially into an archaeal system ancestor of the Tad/Archaeal-T4P and a bacterial system ancestor of the T4aP/T4bP/T2SS/MSH/ComM.

**Fig 3.**
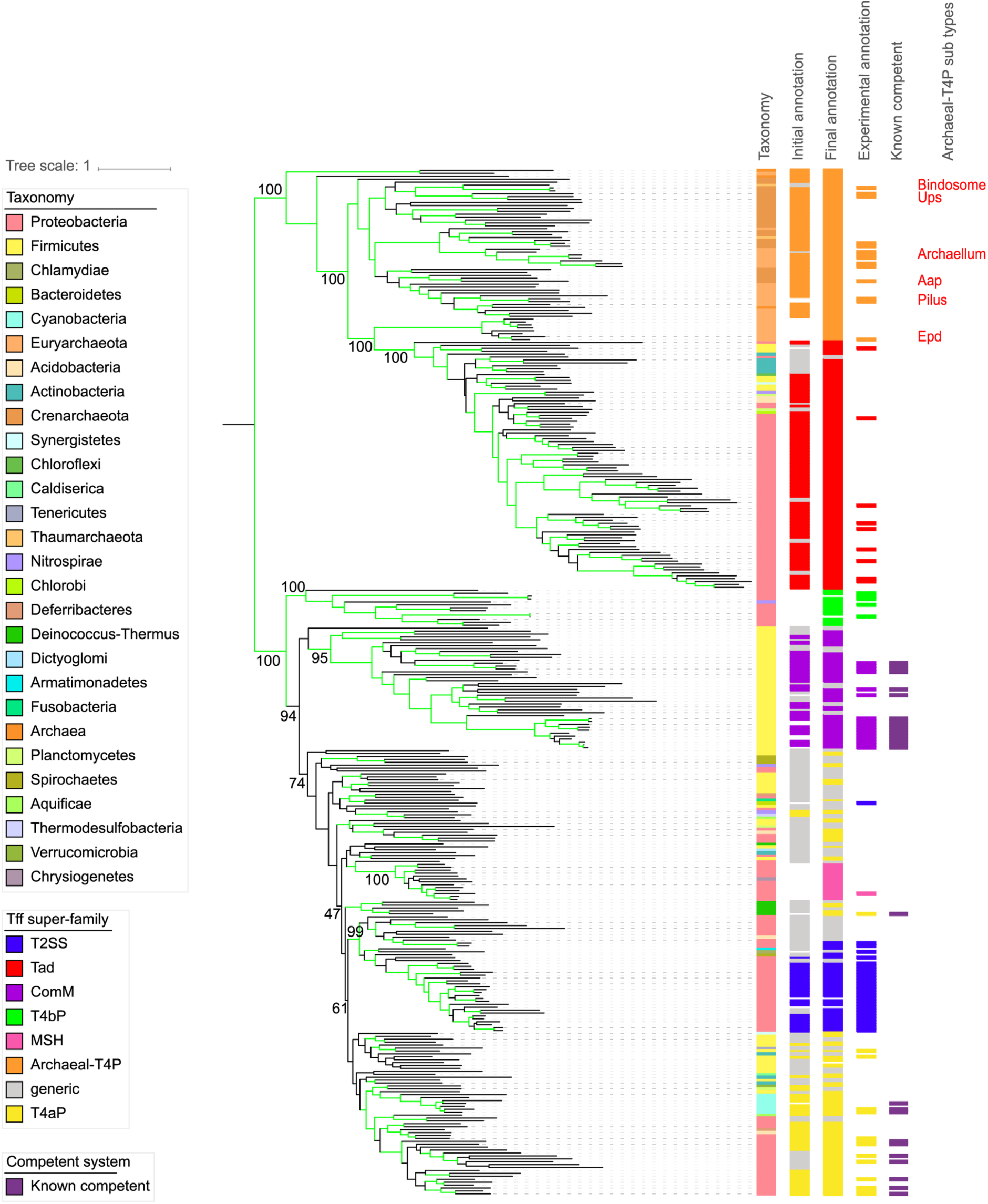
Rooted phylogeny of the TFF-SF. The tree was built with the concatenate of the IM platform (using TadC) and the AAA+ ATPase (using PilB). The colour of the label of the leaves indicates the taxonomic group of the species. The branches are in green if the Ultra-Fast bootstrap is >95%. The supports of the significant nodes are indicated in text. The different coloured strips indicate the classification of the systems with the MacSyFinder annotation (with the initial model and with the final one) and the annotation of the systems in the literature. The systems known to be implicated in natural transformation are indicated in dark purple. Known sub-types of Archaeal-T4P are indicate by a text in red. The tree was built using IQ-Tree, 10000 replicates of UF-Boot, with a partition model.

### The archaeal systems and the emergence of Tad

The ATPase, IM platform and concatenate trees are broadly consistent with five groups within Archaea (Fig. 3, S3, S6, S10), of which four replicate previous findings [37]. All experimentally validated archaella are part of a highly-supported clade (100% UF-boot, group 3 in [37]) that is the sister clade to another highly supported clade containing two pili involved in surface adhesion in Halobacteria (group 2 in [37]). They are sister-groups of a clade gathering the Aap, Bindosome, and Ups (pili described in *Sulfolobus*), as well as other non-characterized pili from Crenarchaeota and Thaumarchaeaota (group 4 in [37]). The rooted tree shows that the basal clade of Archaeal-T4P includes systems mostly found in methanogens (group 1 from [37]). The systems at the base of the tree make a distinct clade but have unknown functions.

Unexpectedly, the position of the root places Tad as a system derived from Archaeal-T4P systems. This feature is found in the trees of the three key components with high confidence. Furthermore, all these trees showed a monophyletic clade including the Tad and the Epd pilus (clade “Epd-like”), whose major pilins have similarly short sequence lengths when compared to the others from Archaeal-T4P (Fig. S7). Both Epd-like pili and Tad have two homologous genes encoding the IM platform, suggesting that their common ancestor already contained them both. We examined the domain structure of these two genes and found that each has one “T2SSF” domain, where most other Archaeal-T4Ps have two such domains and longer IM platform proteins. This strongly suggests that TadB and TadC were derived from an ancestral event of gene fission. To confirm this observation, we aligned the TadB and TadC profiles with the archaeal IM platforms containing two T2SSF domains. In these cases, TadC aligned best with the N-terminal domain, while TadB aligned best with the C-terminal domain of the archaeal proteins. This further supports the gene fission scenario. Finally, the Tad systems have a protein – TadZ – which has significant HMM-HMM profile alignments with Archaeal-T4P components (arCOG00589 and arCOG05608) including those from the Epd-like clade (group 1 from [37]), but not with profiles from the bacterial systems. Altogether, these results strongly suggest that an ancestral Archaeal-T4P harbouring two genes encoding the IM platform diversified into Epd-like systems in Archaea and was transferred horizontally, apparently only once, to Bacteria, leading to the extant Tad systems.

The transfer of the system from Archaea to Bacteria was very ancient. Tad systems were frequently transferred among Bacteria since then (see below), and it is not possible to infer the precise bacterial taxa that acquired the original system. However, the Tad systems at the basis of the clade are from Proteobacteria in 18 out of 20 concatenate trees, often with very good support (Table S3). The two odd concatenate trees place Firmicutes at the base of the Tad clade, but with very low support. This suggests that the ancestor of the Tad system was acquired by a diderm bacterium, and the accretion of the outer-membrane, pore-forming secretin to the Tad system may have been the founding event of these systems. This would fit with the few descriptions of Epd systems that highlight functional similarities with Tad systems [60]. Interestingly, it has been shown that TadD is essential to the assembly of the Tad secretin in *Aggregatibacter actinomycetemcomitans* [61]. While it was originally thought that TadD had no homologs in other systems of the TFF-SF, we observed that it has a homolog in MSH systems (MshN, Fig. 2). It is tempting to speculate that co-option of this component from MSH was determinant to the recruitment of the secretin in the ancestral Tad system arriving at a bacterial diderm.

### The diversification of the bacterial TFF-SF

The other major clade of the TFF-SF only has bacterial systems (T4aP, T4bP, ComM, MSH, T2SS). The vast majority of the concatenate and component trees place T4bP at the basal position in the clade (in the others some generic systems take this position). This is followed by a split between ComM on one side and the T4aP, MSH and T2SS on the other. The T4aP are polyphyletic in all the phylogenetic reconstructions with experimentally validated systems often clustering in the tree (Fig. 3). Some of these systems are in monoderms like Firmicutes and Actinobacteria. The MSH and the T2SS are both clearly distinct and derived from the T4aP. The MSH system falls in a highly-supported clade (100% UF-boot) with other systems of very similar gene composition. T2SS show two exceptions to monophyly. First, the position of Chlamydial T2SS systems next to the other T2SS is highly supported in the ATPase and in the concatenate tree (> 95% UF-boot), but not in the trees of the secretin, major pilin and IM platform. This suggests a chimeric origin for this system where different components were recruited from different types of systems. This may result from the impact of the peculiar developmental cycle and intracellular lifestyle of *Chlamydia* on its envelope [62]. Second, the so-called T2SS of Bacteroidetes (represented by *Cytophaga,* [63]) always cluster away from the remaining T2SS, most often with T4aP, suggesting that they emerged as specialized protein secretion systems independently from the other T2SS.

The key early event in the ATPase trees of the bacteria-only T4P large clade was the amplification leading to the paralogs PilB (the assembly ATPase) and PilT (the retraction ATPase). This event appears as a simple duplication at the base of the tree in certain of the ATPase trees, but also shows more complex scenarios in others (Table S3). In the PilB part of the ATPase tree, the T4bP is basal and the other systems are regrouped with T4aP. This scenario is consistent with that of the secretin tree, where if one places the root between T4aP and T4bP one finds the T2SS deriving from a T4aP system, as in the PilB trees. This is also sustained, albeit with low support, by the major pilin tree, where one finds at basal positions the T4aP and the T4bP. The presence of PilT in very early parts of the tree suggests that the most ancient systems were able to retract the pilus. This is consistent with the ability of both T4aP and T4bP systems to promote twitching motility.

One of the most interesting functions of the super-family, from the evolutionary point of view, is the involvement of some of its systems in natural transformation. The ComM system is commonly found in Firmicutes, even if it is unclear whether it is always involved in transformation. It is monophyletic in all the phylogenetic reconstructions we made, usually with very high support (≥95%). In the concatenate trees, ComM branches apart from a group gathering T4aP, MSH and T2SS, after the divergence with T4bP. The trees of individual components show similar scenarios once one accounts for the effects of the ATPase paralogs, and for the low support of some parts of the IM platform trees. Interestingly, the major pilin trees show ComM branching within T4aP, with poor resolution, close to systems that are known to be involved in natural transformation. This evidence is weak, but suggests a link between the major pilin and transformation. In summary, these results suggest that ComM arose early and only once in the history of the TFF-SF. The T4P systems experimentally linked to natural transformation in diderms were systematically identified as T4aP, and also tend to cluster together in the tree.

### TFF-SF elements are ubiquitous in the prokaryotic world

We used the rooted concatenate tree to class the numerous generic systems that we had previously identified. We assumed that clades where all systems were either generic or of a single type (of which at least one validated experimentally) could be tentatively assigned to that type. Generic systems in clades lacking experimentally validated systems were left unassigned. Only two types of systems were paraphyletic in the tree – T4aP and Archaeal-T4P – and were thus treated differently. T4aP were split in a few monophyletic clades, and systems within each clade were re-assigned using the method above. The Archaeal-T4P systems, from which the Tad derives, can be easily distinguished from the latter, and thus re-assigned, using a taxonomic criterion. This analysis significantly clarified the systems’ assignment (compare Fig S2 with Fig. S11): 1795 out of the 2031 generic systems were re-assigned to classical systems, mostly T2SS (479) and T4aP (748).

We used these tentatively assigned systems to produce more sensitive MacSyFinder models. First, we changed the HMM profiles to account for the genetic diversity introduced by the re-assigned systems. Second, we created models to detect the T4bP and the MSH pilus, since we now had a much larger number of examples of these systems. Finally, we searched for genes systematically associated with the systems’ loci, in a neighbourhood of ±20 genes, that were not matched by any of the HMM profiles of the models. We clustered the proteins by sequence similarity and analysed the largest families. This “guilt-by-association” approach failed to show other proteins systematically associated with a particular type of system (Table S5), suggesting that our models already encompass their most frequent components. This process resulted in more sensitive models that accounted for all known types of systems and correctly identified the 81 experimentally validated systems of bacteria analysed in Table S2, except the T2SS of Chlamydia and Bacteroidetes (shown above to be peculiar).

Using the novel improved models, we found 9026 systems within 4610 genomes, including 1728 T2SS, 2021 Tad, 2558 T4P, 908 ComM, 559 Archaeal-T4P, 177 T4bP, 191 MSH and 884 generic systems (Fig. 4, Table S6). A few systems classed in a given type with the initial conservative models – 14 T2SS, 10 T4P, 5 Tad, 1 ComM – are classed as generic with the new models. However, the inverse is much more frequent, since we re-classified 1114 generic systems as: 1408 T4aP, 338 T2SS, 670 Tad, 4 ComM, 226 Archaeal-T4P. The large number of generic systems re-assigned to T4aP is not surprising, since these systems are encoded in multiple loci, are very diverse and are present in several clades in the tree. This makes them harder to detect using the initial model. The many reassignments of generic systems as T2SS reflects *a posteriori* the excessive stringency of our initial model (based on existing knowledge of systems in Proteobacteria), and the existence of these systems in clades for which there is little or no experimental evidence. The reassignment led to identification of T2SS in a much broader set of taxa including Armatimonadetes, Deferribacteres, Clostridia (presumably diderms, since they include a secretin), Spirochaetes [64], and Aquificae. We also observe many new Tad systems in Elusimicrobia, Actinobacteria, Bacilli and Clostridia (Fig. 4 vs Fig. S2). Our phylogenetics-driven approach for designing new models allowed us to detect diverse putative MSH and T4bP. These systems were so far only described as such in (Gamma) proteobacteria. These are now found for example in Chrysiogenetes and Epsilonproteobacteria for MSH, and in Acidithiobacillia and Nitrospirae for T4bP.

**Fig 4.**
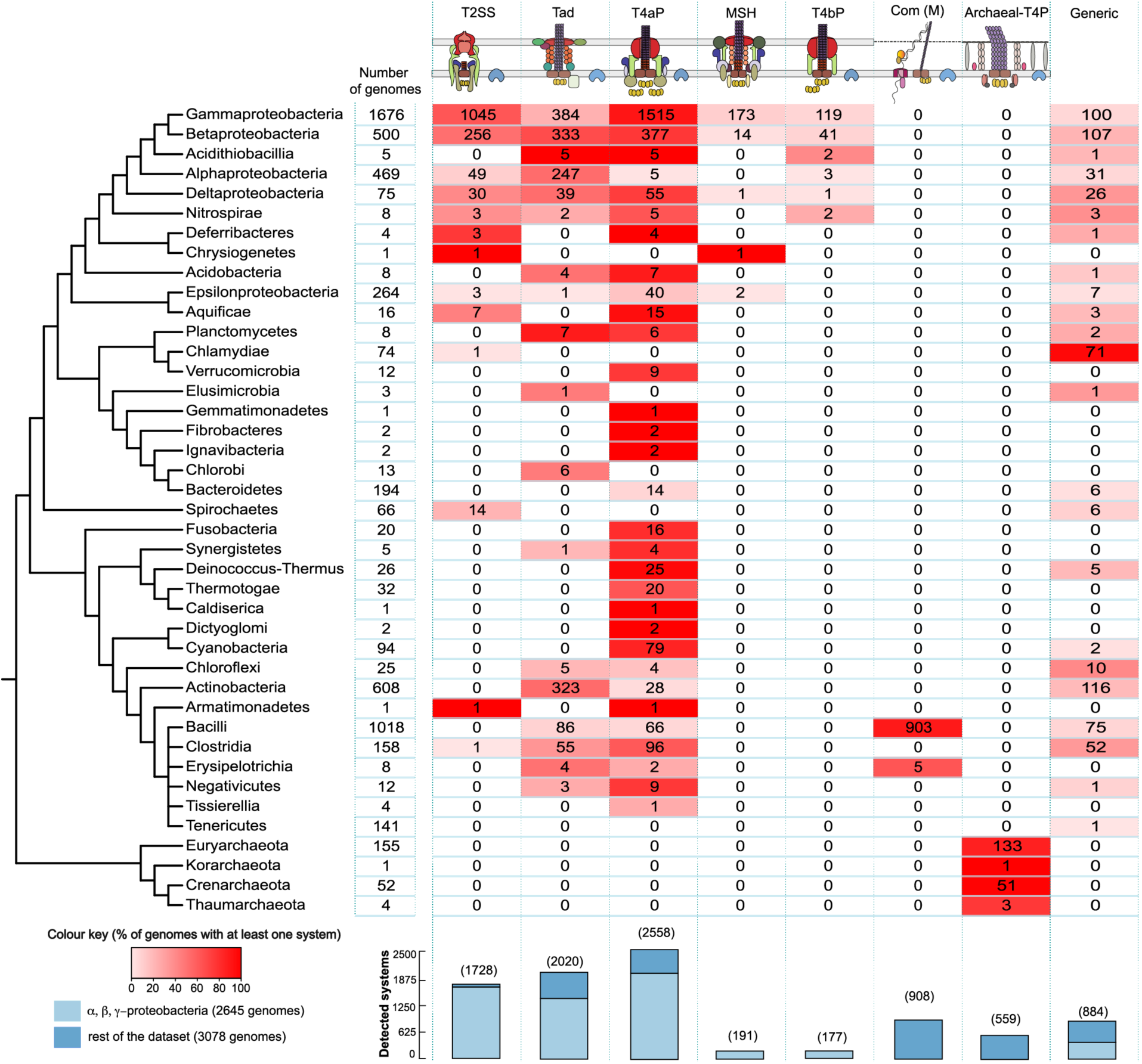
Taxonomic distribution of the systems in Bacteria and Archaea with the final models. Cells indicate the number of genomes with at least one detected system. The cell’s colour gradient represents the proportion of genomes with at least one system in the clade. The bar plot shows the total number of detected systems. The bars are separated in two categories: Alpha-, Beta-, Gamma-proteobacteria versus the other clades. The cladogram symbolizes approximated relationships between the bacterial and archaeal taxa analysed in this study.

In certain cases, the phylogenetic annotation identified some systems that we missed using novel models and this explains the large number of generic systems in certain clades. For example, the new model misses the T2SS in *Chlamydiae* [62] that were annotated with the phylogeny leading to a high number of systems classed as generic in this clade. This is because this system has a small number of components, and we fail systematically to identify members of several families, including the minor pilins and assembly proteins GspLM [62]. Most generic systems in *Chlamydiae* seem to be T2SS given their composition and their location in the concatenate phylogenetic trees. The *Cytophaga* T2SS-like systems could not be identified by the models, which fits the results of the phylogenetic analysis indicating that these are not T2SS *sensu stricto*. Many of the systems of Actinobacteria remain classed as generic systems. A large fraction of them are classed as Tad by proximity to experimentally validated systems in the phylogeny, but they lack identifiable homologs of some usual components such as the minor pilins and TadC (their TadB does not contain two domains like those of homologs in some Archaea, showing this is not the result of a gene fusion). Finally, the Clostridial systems are classed as generic by the models, but phylogenetically they tend to be close to the remaining ComM systems.

In conclusion, the final models classify more systems and assign them classifications that are in most cases consistent with the phylogenetic analysis. Some discrepancies subsist that can be due to systems very divergent from the models (missed by the models) or inactive systems (spuriously picked by the generic model). Remarkably, this refined annotation shows that all represented phyla of Bacteria and Archaea have at least one system from the TFF-SF, underlying the huge evolutionary success of this potentially very ancient type of machinery.

### Genetic organization is associated with differences in rates of horizontal transfer

The systems differ strikingly in terms of genetic organization (Fig. 5). ComM and T4aP are usually found in multiple loci, whereas MSH and Tad are almost exclusively encoded in a single locus. Hence, as systems diverged their genetic organization also changed. To detail the prototypical genetic organizations of each type of system, we built a graph where nodes represent components and edges link components that are encoded contiguously in the genome. The edges are weighted by the frequency of contiguity: genes that are systematically contiguous are linked by thick edges. This graph reveals prevailing genetic organizations for most types of system (Fig. 5, S12). The operon *comG*[*ABCDEFG*] is usually conserved and apart from the other components in ComM. The T4P genes are usually scattered in four operons. The T2SS is typically encoded on a single locus usually in a conserved order, with some exceptions exemplified by the *Legionella* system, and the Tad has a large locus with some elements showing some local permutations of gene order. No clear genetic organization or prototypical repertoire emerged from the contiguity graph built for Archaeal-T4P, which is partly a consequence of the large number of homologous families in the system (Fig. S12), and partly of their diversity in terms of gene content and function. The analysis of the representative Archaella systems shows more conserved genetic organization, even if many variants exist [37, 44] (Fig. S13). Interestingly, the genetic organization of Epd is very similar to the Tad: the two IM platform genes are contiguous and followed by the major ATPase and the secondary one (TadZ in Tad and FlaH (arCOG04148) in Epd) (see Fig. 5). Hence, the core genetic organization of Tad evolved in Archaea before the transfer of the system to Bacteria.

**Fig 5.**
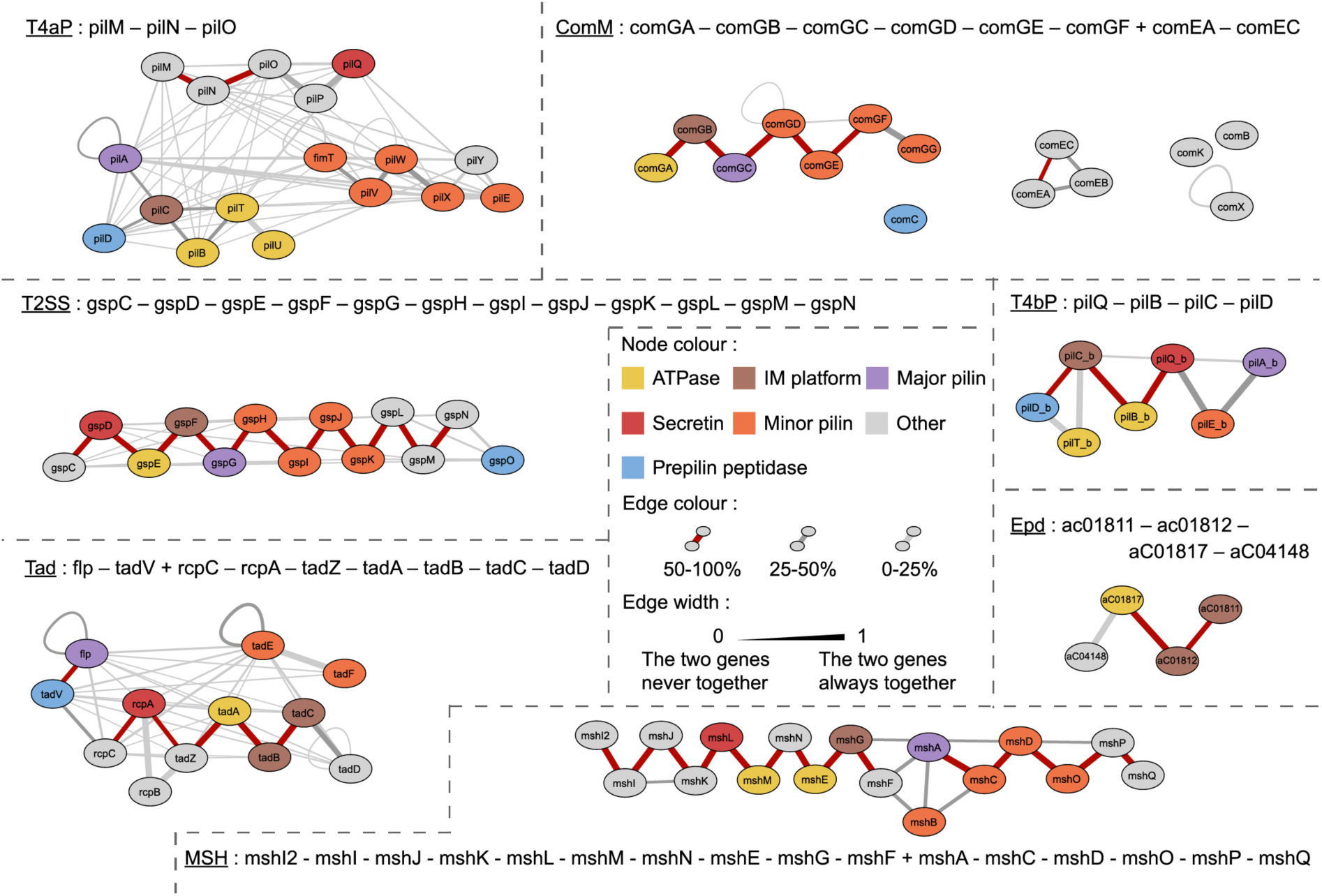
Genetic organization of the detected systems. For each detected system (those indicated in Fig. 4), the edge width represents the number of times the two genes are contiguous divided by the number of times the rarest gene is present in the system. The colour of the edge represents the number of times the two genes are contiguous in the system divided by the number of systems.

The patterns of genetic organization of the homologous components differ between systems. For example, T4aP shows a conserved triplet of genes *pilBCD* of which the pair *pilBC* is often conserved in other types of systems but *pilCD* is not. In general, pilins tend to be encoded together, but can vary in their co-localization with the rest of the genes: they can be apart (T4aP), at the edge of the locus (ComM, Tad) or in the middle (T2SS, T4bP). In archaea, all cases were found. Interestingly, many duplicated genes tend to be contiguous. This is the case of many pilins, of the *pilUT* genes encoding the ATPases, and of the integral membrane platform genes *tadBC*. This is consistent with models suggesting a bias towards gene duplication in tandem [65]. The variability between types of systems and the conservation within types, suggest that genetic organization is under selection within types, but changes rapidly upon functional innovation.

The genetic organization of the loci can also reflect the action of horizontal gene transfer. If the systems are often gained or lost within lineages, as it was shown for Tad [66] but much less so for the Achaellum [44], then systems encoded in a single locus are much more likely to be successfully transferred because all the necessary genetic information can be transferred in one event [67]. Systems scattered in different distant loci cannot be transferred in a single event (although parts of the system can presumably be exchanged if the recipient genome encodes the system). We thus hypothesized that systems encoded in single loci are more likely to undergo horizontal gene transfer. To test this hypothesis, we compared the phylogenetic tree of each system, a sub-tree of the larger phylogenetic reconstruction, with a maximum likelihood tree of the 16S rRNA sequences of the species carrying the systems (Fig. S14). We excluded the archaeal systems from these analyses because their loci are harder to define precisely (sometimes scattered and multiple systems per genome) and their functions are still poorly delimited in most cases (complicating the definition of the clade to use in the analysis). We found that systems encoded systematically in a single locus are more frequently transferred than those encoded in several loci (Fig. 6). These results are reinforced by the analysis of the frequency with which systems are encoded in plasmids, which follows closely the trends observed for the frequency of transfer (highest in Tad and lowest in ComM, Fig. 6). The contrast is especially interesting between the Tad and T4aP systems that are both present in many different clades and are encoded almost exclusively in one locus (Tad) or many loci (T4aP). This association between rates of transfer and organization suggests that systems that are frequently gained and lost endure a selective pressure for being encoded in a single locus.

**Fig 6.**
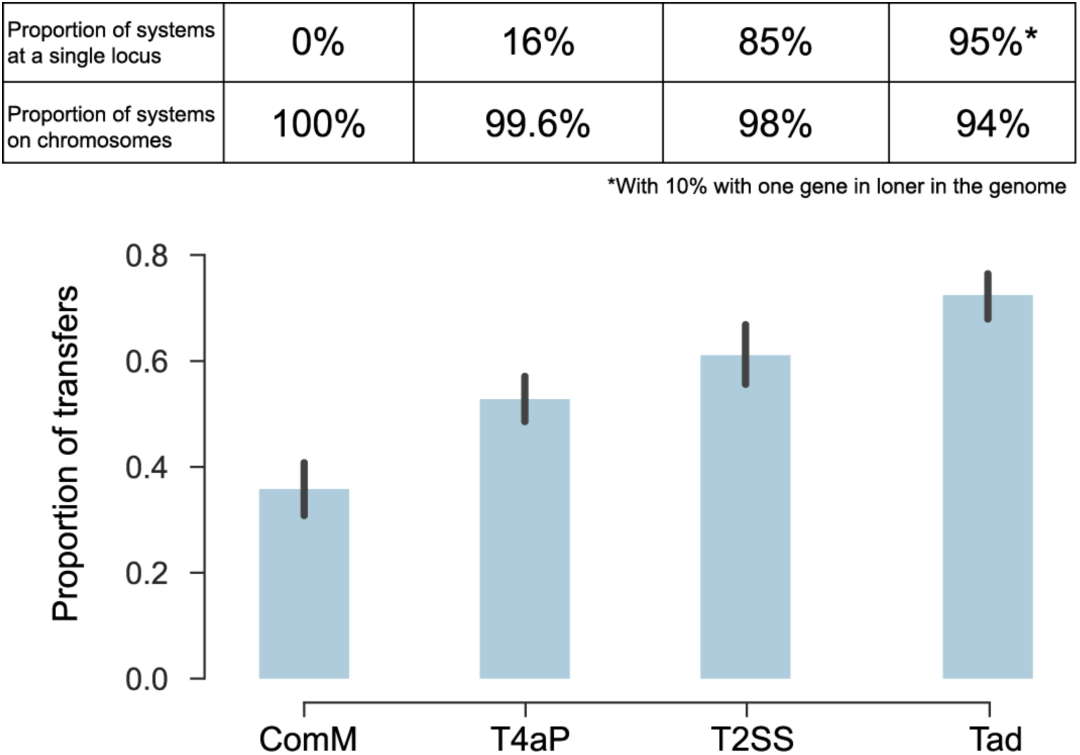
Association between organization and horizontal transfer of the different systems. For each system, we compared the subtree of the systems with the 16S tree of the same species using ALE v0.4 [113] to obtain the proportion of transfers. We also show the proportion of systems in single locus and the proportion of systems on chromosomes (the others being found on plasmids).

## DISCUSSION

We used comparative genomics and phylogenetics to produce models and protein profiles that identify TFF-related systems in genomic data of all Prokaryotes. They are publicly available and provide a significant advance relative to our previous work, since they are more sensitive and cover more types of systems (Archaeal-T4P, ComM, MSH and T4bP). We used them to quantify the frequency and taxonomic distribution of the different systems. Strikingly, every inspected phylum of Prokaryotes has some type, and often several types, of systems from the TFF-SF. Some types are more widespread (e.g. T4aP, Tad) than others within the limitations of the taxonomic coverage provided by current genome databases. Notably, the MSH and T4bP, for which few experimentally validated systems are known, are abundant in Proteobacteria, but absent from most other phyla. For the archaeal systems, most clades outside of archaella are associated to a very limited number of characterized systems with distinct functions, highlighting the large genetic and functional diversity of Archaeal-T4P. Further experimental study of these systems is required to produce reliable MacSyFinder models for each of them.

Our approach may be regarded as conservative. First, some components of the systems are not sufficiently conserved in sequence for reliable phylogenetic analyses at this large time-scale and were not used in the phylogenetic inference. The minor pilins are a particularly important set of proteins that were ignored because they produced short and very poor multiple alignments. Second, we rely on the existence of experimentally validated systems and on monophyletic clades having such systems to build the models. If the systems have been described in few species, or in a small number of phyla, then this limits our ability to identify them, especially when they are very different in terms of gene repertoires and protein sequences. This may explain why our models missed the T2SS of Chlamydiae: they carry few components and these are of different origins. Notably, its major pilin is atypical with a disulphide bond and presumably lacking the Ca-binding site. In other cases, systems may actually differ from the descriptions in the literature. This is probably the case of the so-called T2SS of Bacteroidetes. This system is involved in protein secretion [63], but consistently branches apart from T2SS in all analysis of the phylogenetic markers. The major pilin of this system is very divergent compared to major pseudopilins from proteobacteria. Our analysis raises the exciting possibility that it might represent a novel type of secretion system derived from the T4aP independently of the T2SS. Actually, all trees show that the widely studied T4aP systems are very diverse and form several different clades in the tree, where the one with the PilU ATPase, the most widely studied, accounts for a minority of the identified systems. Most of the other T4aP are poorly characterised and may represent systems with novel properties. Finally, the results obtained with the final improved models showed few systems identified as generic (once we exclude the systems resembling T2SS in Chlamydiae, the ComM in Clostridia and the Tad in Actinobacteria). This suggests that there may be few, if any, radically novel systems to be discovered in the super-family that contain the three key components (ATPase, IM platform, major pilin). On the other hand, the diversity of certain types of systems – like the T4aP – may still reveal surprising novel functionalities.

The phylogeny of the key components of the TFF-SF revealed an initial split between archaeal and bacterial systems, suggesting that these structures may have pre-dated the last common ancestor of all cellular organisms (Fig. 7). This ancestral system presumably had one ATPase for its assembly (the function performed by PilB in T4aP), an integral membrane platform, pilins and a prepilin peptidase. Among these key components, only the ATPase has identifiable sequence homologs outside the super-family. The PFAM domain of the prepilin peptidase of T4P belongs to the PFAM clan CL0130 with other signal-peptide inner membrane-associated peptidases, several of which are found in Bacteria, Eukaryotes (the presenillin family proteases), and Archaea [68]. However, we cannot exclude that other protease(s) might perform this function. Hence, an enzyme able to mature pilins should already have been available in the organism encoding the ancestral system. The protein profiles of the integral membrane platform match those of some ABC transporters, and the protein is structurally very similar to one of the V-type ATP synthase subunits [69]. This suggests that the original component may have been co-opted from these ubiquitous membrane-associated systems. The pilins are small and evolve fast; it is impossible to trace their evolution using sequence analysis at deep time scales. Fast evolution of pilin globular domains may be associated with the variability of essential inner membrane components that promote pilin targeting to the assembly site, or connect the inner and the outer membrane sub-complexes [70, 71]. It is also difficult to detail if there were other components in the ancestral system of the super-family, since they either evolve fast or are present in only a small number of systems. A recent study showed that a minimal set of eight genes was sufficient to produce assembly of the T4aP of *Neisseria meningitidis* [72]. Four of them, PliMNOP, are essential for the assembly but are lacking in our list of ancestral genes because their homologs could not be found in any or in the vast majority of genes neighbouring T4bP, ComM, Tad and Archaeal-T4P. They were found in MSH and T2SS (Fig. 2, Table S5), suggesting that they arose more recently and that other systems do not require these proteins for assembly (Fig. 7). In short, our results are consistent with the idea that the ancestral system was able to energise its assembly and build up a pilus with matured pilins on top of an assembly platform, the basic molecular architecture of all extant systems.

**Fig 7.**
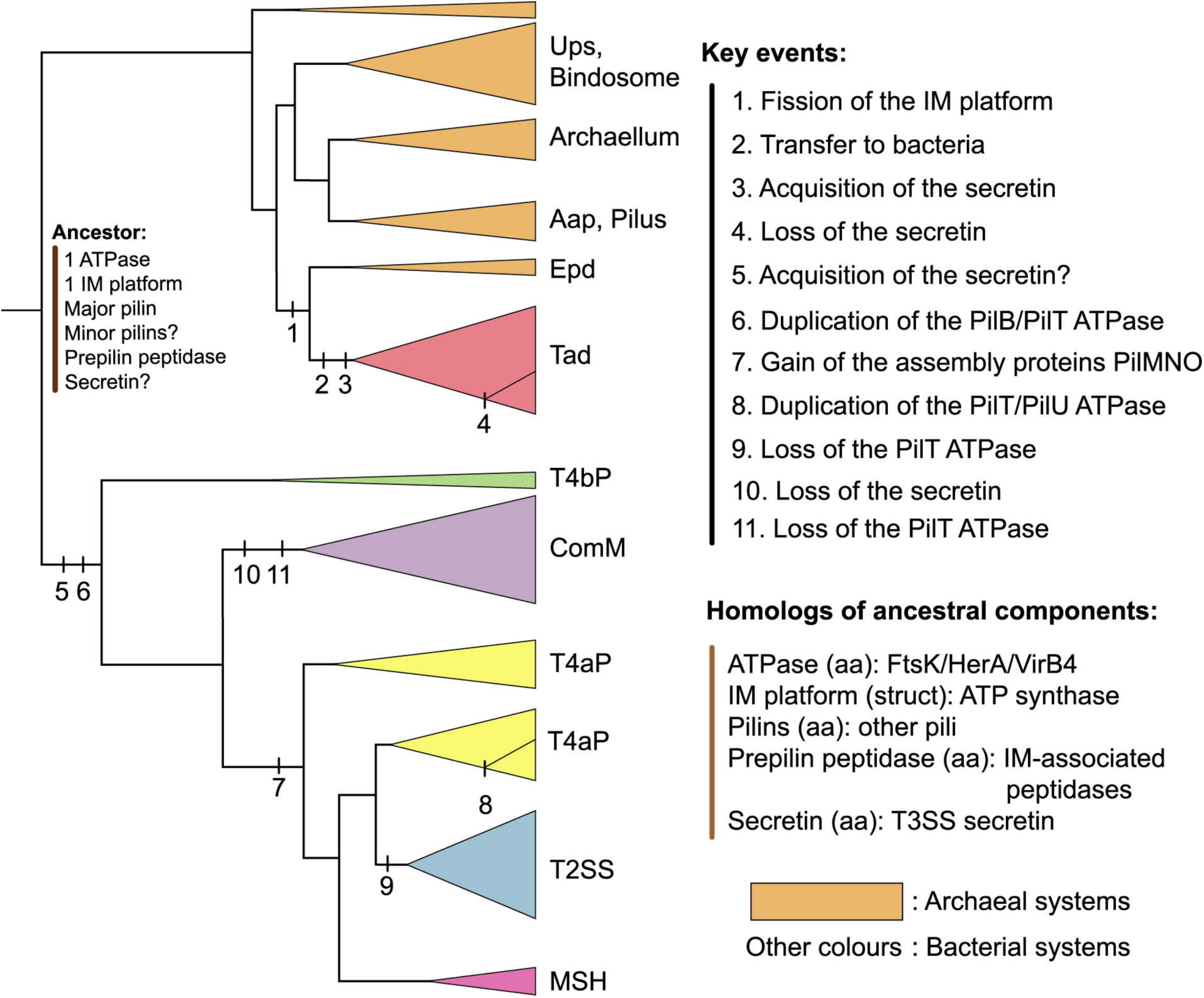
Evolutionary scenario of the TFF-SF. The tree was based on the information of the trees of the concatenate and simplified to highlight the key clades and events. The colour of the triangles indicates the type of the systems. Each vertical bar on the branch indicates a numbered potential evolutionary event, whose details are specified under the corresponding number in the list “Key events”. The hypotheses for the composition of the last common ancestor of the TFF-SF are indicated at the root, and the distant homologs of these systems are indicated in the list “Homologous of ancestral components”, where homology was observed by sequence (“aa”) or structural (“struct”) similarity.

Ours and previous data on the genetic composition and organization of archaeal systems [37], reveal processes of functional diversification leading to several families, tentatively associated with different functions, including a system – precursor of the Tad and resembling extant Epd systems– that was transferred to Bacteria. Archaeal-T4P are present in many clades of Archaea. The *Sulfolobus* genus alone counts systems from four of the seven different archaeal-T4P types studied experimentally (Aap, Bindosome, Ups, and archaellum). Even though horizontal transfers might be frequent among Archaea, our approach places the root of Archaeal-T4P within systems of methanogens from the Euryarchaeota phylum (group 1 of Makarova et. al [37]) – consistent with a proposed rooting for the archaeal tree of life within methanogens [73]. Only performing experiments on more diverse systems and species would enable to elucidate the wide range of functions of the Archaeal-T4P.

The Epd-like systems share the closest ancestry with Tad systems among all elements of the TFF-SF. They were only characterized in *Methanococcus maripaludis*, where they are involved in surface attachment, a trait they share with the Tad pilus [60]. The Archaeal origin of Tad is consistently suggested by the rooted phylogenetic analyses and the specific shared characteristics of pilins, IM platform and TadZ-like proteins in Tad and archaeal-T4P. The literature often classes Tad pilus as T4bP [74]. Our study shows that these systems are very different in terms of components and genetic organization and distantly related from the evolutionary point of view. This is in accordance with recent works proposing to clearly separate Tad from T4bP and to name them as T4cP [45]. A striking trait of Tad (and Epd) is the systematic presence of two genes (*tadB* and *tadC*) encoding the IM platform. This has been regarded as the result of a gene duplication [39], but the size and domain content of these genes is more parsimoniously explained by a gene fission event, e.g. by a mutation integrating a stop codon within the ancestral gene. This produces a complex evolutionary scenario: the original IM platform has two homologous domains, suggesting a duplication before the last common ancestor of the TFF-SF. In the Epd and Tad clades this was followed by a fission event. The adaptive relevance of these successive events in the light of emerging structural data could be an interesting topic of future research.

The concatenate tree suggests that Tad’s ancestor was transferred from Archaea to a diderm Bacteria where it acquired a secretin and then diversified and transferred throughout the bacterial domain. The secretin tree provides some information of when the initial transfer took place. This tree places Tad’s secretin within those of T4aP systems with high confidence and typically close to Proteobacteria. The functions of this ancestral system should resemble the ones of extant ones, since the closely related Epd systems in Archaea have similar genetic organization (this work) and are also involved in similar functions [60]. This suggests that the co-option of the secretin upon transfer of the ancestor to a diderm was the founding event of Tad systems. It occurred at a time when most types of systems (T4aP, T4bP, ComM, and possibly MSH and T2SS) were already in place. This hypothesis is reinforced by the secretin tree showing one single monophyletic clade for Tad’s secretin, suggesting that accretion of the secretin to this system only happened once. Co-options of a secretin from other systems are very common. They were observed multiple times in the evolution of T3SS (e.g., from Tad and from T2SS) and in filamentous phages [11]. If this scenario is correct, then the adaptation of Tad to monoderms, which occurs at several places independently in the tree, involved the loss of the secretin. This event of loss seems very common since it is also found once in the initial evolution of ComM and several times at the emergence of T4aP of monoderms. Finally, the widespread distribution of Tad, in spite of its relatively recent origin is in agreement with the very high observed frequency of horizontal transfer for this system.

The T4bP is the most basal system among Bacteria. T4aP and T4bP are usually distinguished based on their molecular size and sequences of their leader signal peptides [75]. The T4aP constitutes a thin pilus (5-8nm diameter), whereas the T4bP pilins are longer [76, 77]. The peptide signal of T4bP pilins tends to be larger than those of the T4aP [78]. These differences match the early divergence observed for T4aP and T4bP. Subsequently, a split separated ComM from T4aP, and the latter then diversified into T2SS and MSH. Recent works suggest that the last common ancestor of Bacteria was a diderm [79]. Our analysis shows that Tad, T2SS and T4aP are monophyletic clades in the phylogeny of the secretin, in agreement with previous works [80], suggesting that there was little transfer of the secretin between systems. Furthermore, if one roots the secretin tree between T4bP and T4aP (in agreement with the concatenate tree of the TFF-SF), then the tree of the secretin largely recapitulates the tree of the concatenate of the super-family (except for Tad). This is in line with a very ancient acquisition of the secretin by the TFF-SF. Since T4bP are the most basal systems in the tree and are only found in diderms, this strongly suggests that the original bacterial system had a secretin and was present in a diderm.

Until recently, it was thought that only systems encoding PilT were capable of pilus retraction. This would suggest that the ability to retract the pilus resulted from the neo-functionalization of one of the copies (PilB in T4aP) of the protein at the moment of the duplication of the ATPase leading to PilB/PilT in T4P. Surprisingly, it was recently shown that the Tad system of *Caulobacter crescentus* is also able to retract the pilus [45]. As these authors, we only identified one copy of the ATPase in this system, in agreement with the data that PilB can also retract the pilus. It is thus possible that the original ATPase of the ancestor system of T4bP and T4aP could perform both activities – assembly and retraction/disassembly – and that the duplication led to the sub-functionalization of these functions. In support of this model, the T4aP in *Vibrio cholerae* was shown to retract with low speed in the absence of PilT [81]. Interestingly, even though not demonstrated, it was previously suggested that Archaeal-T4P retraction might be required to explain some archaeal communities’ behaviour, like transition from sessile to swimming stages [82]. Overall, there is now a large set of results pointing to the ability of a single ATPase (PilB in T4aP) to make extension and retraction of the pili. Some overlap between the functions of PilT and PilB may also explain why PilT is frequently lost (e.g. in T2SS and ComM). It is tempting to speculate that other independent duplications of PilT/PilB scattered in the phylogenetic tree represent further specialization of the activities of the ATPases.

The increase in the diversity of systems able to retract the pilus opens the possibility that many other pili could be involved in natural transformation, since the pilus role is to attach the DNA and bring it to the cell surface. Actually, a number of arguments are in favour of the hypothesis that the ancestor of the bacterial systems might have been able to facilitate natural transformation. First, ComM, T4bP and T4aP have all been associated with this mechanism and they are among the most basal bacterial systems in the tree. Second, the predicted repertoire of genes of the last common ancestor of these types of systems could suffice for DNA attachment and retraction towards the cell envelope necessary for transformation. Third, systems that emerged within Archaea are able to facilitate transformation. The Ups pilus in *Sulfolobus* is highly expressed under UV light, mediates cell aggregation and facilitates natural transformation mediated by the independent *ced* system [83, 84]. A Tad locus (Archaea-derived like all Tad) from *Micrococcus luteus* has recently been shown to be required for natural transformation [85]. We identified this system within the Tad clade and its gene repertoire includes one single ATPase. Fourth, it has been previously shown that other key components of the transformation machinery – DprA and ComEC – are widespread across Bacteria [34, 86]. It has been known that the DNA uptake machinery required for transformation is encoded in many bacteria that were never shown to be naturally transformable [87, 88]. For example, *Escherichia coli* and other enterobacteria contain functional T4aP genes coregulated with the competence machinery [70, 89], which are required for natural transformation [90]. If many of the elements of the TFF-SF can pair with this machinery to make a cell naturally transformable, then a vast majority of Bacteria could potentially be naturally transformable. Interestingly ComM and T4aP systems known to be involved in natural transformation tend to cluster in the phylogenetic tree of the concatenate. This suggests that even though many T4P might facilitate transformation, those effectively involved in transformation have evolved certain traits improving this function. One such feature is the presence of two disulphide bonds, which may stabilize the structure and improve retraction force-resistance of *Acinetobacter* [91] and enterobacterial T4aP major pilins [70], or of the competence-specific minor pilins in Neisseria [92].

Our study has revealed how a small set of proteins with different functions evolved to produce different adaptive functions involved in different types of motility, adherence, DNA acquisition and protein secretion. This process involved: 1) Accretion of accessory proteins, such as the secretin and secretin-associated proteins to cope with the existence of an outer membrane in diderms. 2) Duplication and sub-functionalization of key components, such as pilins and ATPases. 3) Gene fission in TadBC. 4) Several cases of gene loss, notably for the IM platform paralogs in Tad and some archaeal systems, for the PilT ATPase, and for the secretin in monoderms. 5) Gene transfer between distant clades, including a rare example of a large macromolecular system (Tad) transferred from Archaea to Bacteria. 6) These events were accompanied by rearrangements of the genetic loci. TFF-SF systems were frequently transferred horizontally, which certainly accelerated their evolution, since genetic exchanges break clonal interference and accelerate innovation processes by recombination [93]. Interestingly, we observed that genetic organization and horizontal transfer were intimately associated, with systems encoded in one single locus showing higher rates of transfer. This may be a general pattern in the molecular evolution of complex systems in Prokaryotes. Strong genetic linkage facilitates positive selection in physically interacting proteins [94], and facilitates the spread of the system to other species [67]. Novel genetic contexts may in turn select for further changes in the systems. Once functions remain a long period of time in the lineage, as seems to be the case for ComM and some T4aP, major adaptive changes in the systems may become rare and rearrangements splitting the initial locus may be eventually fixed. Radical changes among split loci are less likely to be spread by horizontal transfer, unless the recipient cell has already a copy of the system with similar genetic organisation. As a result, a tight association is established between genetic organization and the ability of a system to evolve and spread by the action of horizontal gene transfer.

## Methods

### Data

We analysed 5768 complete bacterial and archaeal genomes from NCBI RefSeq (ftp://ftp.ncbi.nlm.nih.gov/genomes/refseq/, last accessed in November 2016), representing 2268 species of Bacteria and 168 species of Archaea.

### Detection of the TFF super-family

All the systems of the family were detected using MacSyFinder v1.0.2 [46]. This program uses a model to identify a type of system in a DNA sequence (typically a replicon). The model specifies the components of the system, each represented by a HMM profile, and how their systems are organized in the sequence. A full description of the program and the models can be found in: http://macsyfinder.readthedocs.io/en/latest/index.html. Briefly, the components can be mandatory, accessory, or forbidden. A system is only validated if it fulfils a quorum of mandatory (MMGR: minimum mandatory genes required) and/or mandatory + accessory genes (MGR: minimum genes required). A locus is excluded if it contains a forbidden gene (these are useful to discriminate between closely related systems with a few specific components). Components are expected to be clustered in the genome at a short distance (defined in the model). Yet, some components can be defined as “loner” and be encoded apart. A component can be set as “exchangeable”, in which case several HMM profiles can be used to detect it (for example the same prepilin peptidase is used by T2SS and T4P in some cases [52–54], and both profiles can be used to identify the prepilin peptidase of each of the two types of systems).

For this work, we could use the models previously proposed by TXSScan [41] for T2SS, T4P, and Tad, but we wished to add a few components that were missing there. For the Archaeal-T4P and for ComM we did not have an initial model. We proceeded in two steps. First, we made conservative *initial model*s that matched the archetypal systems, but sometimes were too strict for some atypical systems. This resulted in a list of systems in which we had strong confidence. However, it also missed many systems. To identify these systems, we built a model called *generic* that had only the basic building blocks of these systems, with all the homologous proteins set as “exchangeable”. Following the comparative and phylogenetic analyses we re-defined all the models to make *less-initial models* that could identify a larger number of systems. Both sets of models are made available in Dataset S1. The table with all protein profiles is given in Table S4.

#### Generic

We defined the model called *generic* to search for variants of the TFF-SF that include the key components but do not fit the strict definitions of the more specific models (T4aP, T2SS, etc). This model assumes that all the HMM profiles of the same connected component in the profile-profile graph of similarity can fill in for the function. Hence, it can identify very divergent or minimalistic systems as well as chimeric systems with components that match profiles from different types. A cluster of components is classed as generic if it does not fit any of the more specific models and contains an ATPase, an integral membrane platform, and a major pilin. In addition to these three proteins, the generic model also includes the prepilin peptidase and the secretin that are not deemed essential for the system because the former may be recruited from other systems in the genome [54] and the latter is specific to diderms.

#### Tad

The *initial model* of Tad followed closely the definitions proposed in [41]. This model includes all the known key components of the system and assumes that they are all encoded together, with the exception of *tadV,* the prepilin peptidase gene that can be encoded apart (loner) and be exchangeable with a number of homologous components from the T4P (*pilD*) and ComM (*comC*).

The *final model* includes *tadD, rcpB* and *rcpC* as new accessory components. The prepilin peptidase TadV is no longer exchangeable. The model defines the Tad pilus as multi_loci to allow for the existence of systems encoded in loci scattered in the genome (even if this is very rare).

#### T4aP

The *initial model* of T4aP was significantly improved from the model in [41]. It is more precise in the annotation of the retraction ATPases (*pilT* et *pilU*), the major (*pilA*) and minor pilins (*pilE*, *pilX* and *fimT*), accounting for five novel components: *pilT, pilE, pilA and fimT* set as mandatory, and *pilU* and *pilX* set as accessory, according to their occurrence in the systems. Accordingly, the number of MGR and MMGR were increased to 8. The prepilin peptidase *pilD* was changed to mandatory, loner and exchangeable with a number of homologous components from the T2SS (*gspO*) and ComM (*comC*) according to its localization that could be found alone in the genome and the fact that the HMM profile of these two genes often have better e-value than the one of the T4aP.

The *final model* of T4P includes *pilW*, *pilX* and *pilY* as new accessory components. We decreased MMGR to 4 and MGR to 5, which fits better the data. We set *fimT, pilM, pilP* and *pilA* as accessory, to help MacSyFinder to search more complete T4P in the genome. We also removed the forbidden genes *gspN*, *tadZ* and *gspC*.

#### T2SS

The *initial model* of T2SS Tad followed closely the definitions proposed in [41], where we increased the MGR to 8 and set the prepilin peptidase *gspO* as mandatory, loner and exchangeable with a number of homologous components from the T4aP (*pilD*).

The *final model* was relaxed to identify a larger fraction of the systems. We reduced the MMGR to 4 and the MGR to 5. To fit the data better, we added the prepilin peptidase of ComM (*comC)* as another exchangeable gene of *gspO*. We set *gspC* gene as mandatory, *gspM* as accessory and *gspD* was set as loner to fit better the data.

#### ComM

In this *initial model* only the genes that compose the pilus were used in the model without the genes that encode DNA uptake system, such as *comEA*, *comEB* and *comEC* [34, 95-98]. The minimal distance between genes was set to 5. The MMGR was set to 3 and the MGR to 5. And the system was set as multi_loci as some genes are loner. The genes *comC*, *comGA*, *comGB*, *comGC* and *comGD* were set as mandatory and the other ones were set as accessory in relation with their presence in experimentally validated systems, curated with an exploratory phase to know the relative abundance of the genes in the systems (the genes with more than 80% of presence in the detected systems were set as mandatory and the other as accessory). *comC* was set as loner and exchangeable with *pilD* of the T4aP because we found case where the HMM profile of *pilD* was better in e-value than the *comC*, same for the *comGA* was set as exchangeable with *pilB* of T4aP. The genes *comB*, *comK* and *comX* were set as loner because they are often found alone in the genome.

In the *final model* we changed the number of genes for the MMGR and MGR to 4. We also added the genes encoding the DNA uptake system in the plasma membrane (*comEC*, *comEB*, *comEA*). *comEC* was set as mandatory *and comEB* and *comEA* was set as accessory, and *comEC* and *comEB* were set as loner. Changing the gene *comGD* to accessory allowed us to search for loner gene without changing the MGR number.

#### Archaeal-T4P

*Initial model*. We here describe the first tool to detect archaeal T4P. We extracted the sequences from the 200 arCOG families (2014 version, [99]) deemed to be associated to archaeal T4P by Makarova and colleagues [37]. We built HMM profiles for each of these families: sequences were aligned with MAFFT v7.273, linsi algorithm, and the alignment extremities were trimmed based on the results of BMGE with BLOSUM40 matrix [100, 101]. HMM profiles were generated using Hmmer version 3.1b2 [102]. These profiles were compared to profiles of Tad, T4P, and T2SS from TXSScan [41] using HHsearch (e-value and p-value threshold of 0.001 for the family cutoff), in order to define supra-families of components [48]. Core “mandatory” components were defined based on the literature and experimentally validated systems. Other components were set as “accessory”. The arCOG families that matched the same component were defined as exchangeable. The prepilin peptidase was set as a loner gene that can be part of multiple systems. This initial model asked for a minimal number of mandatory genes and overall number of genes of 4. Of 14 experimentally validated systems found in the literature, 10 were detected with this initial model (Table S1). After counting the occurrence of the different arCOG in the detected Archaeal-T4P, we removed those without any occurrence, to reduce the number to 109 arCOG families.

*Final model*. The number of genes for MMGR and MGR was reduced to 3, which fits better the data.

#### T4bP

*Final model.* This class includes the R64 thin pilus, toxin-coregulated pilus, bundle-forming pilus, longus pilus and Cof pilus [29]. As we do not have many experimentally validated systems for the T4bP, we used the phylogenetic information of the TFF-SF trees to have a set of T4bP-related proteins to create the HMM profiles and the definition of the model. We created 8 HMM profiles, the MMGR was set to 4 and the MGR to 4. The system was set as multi_loci as some genes are loners. The genes *pilD*, *pilB*, *pilA*, *pilC* and *pilQ* were set as mandatory and the other ones were set as accessory, according to their occurrence in the detected systems. The prepilin peptidase *pilD* was set as loner. The *pilA* was set as exchangeable with *pilA* of the T4aP because we found cases where the HMM profile of *pilA* of T4aP had a better e-value in matches to T4bP than the *pilA* of T4bP.

#### MSH

*Final model.* As we do not have many experimentally validated systems for the MSH, we used the phylogenetic information of the TFF-SF system tree to have a set of MSH-related proteins to create the HMM profiles and the definition of the model. We created 20 HMM profiles, the MMGR was set to 3 and the MGR to 4. The system was set as multi_loci as some genes are loner. The genes *mshA*, *mshE*, *mshG*, *mshL* and *mshM* were set as mandatory and the others were set as accessory, according to their occurrence in the detected systems. The gene *mshA* was set as loner, according to detected systems found in the genomes. The *mshB* was set as exchangeable with *pilA* of the T4bP because we found cases where the HMM profile of *pilA* of T4bP provided better e-values when matching MSH systems than the *mshB* of MSH. For similar reasons, *mshC* was set as exchangeable with *fimT* of T4aP.

### Retrieval and construction of protein profiles

We retrieved 37 profiles for T2SS, T4aP and Tad from TXSScan [41]. For two HMM profiles of T4aP that combine the detection of two proteins paralogs (T4P_pilT_pilU and T4P_pilAE), we decided to separate from the original alignment of this profile the sequence of the different proteins to generate five separate HMM profiles (T4P_pilT, T4P_pilU, T4P_pilA, T4P_fimT and T4P_pilE).

To create the HMM profiles, we used the following methodology. For the genes that had few representatives in the experimentally validated dataset, we used BLASTP v 2.5.0+ (default settings, e-value < 10^−6^) [103] to search for homologs among complete genomes. To remove very closely related proteins, we performed an all-against-all BLASTP v2.5.0+ analysis and clustered the proteins with at least 80% sequence similarity using SiLiX v1.2.10-p1 (default settings) [104]. We selected the one sequence from each family as a representative. We aligned all the representatives using MAFFT v7.273 (--auto, automatic selection of the parameters depending of the size of the alignment, default values for the other parameters) [100]. With SEAVIEW [105], the poorly aligned regions at the extremities were manually trimmed in the alignment. The trimmed alignment was used to build the HMM profile using hmmbuild (default parameters) from HMMER package v3.1b2 [106].

For the HMM profiles of the final model, we used the sequences of the profiles describe above. Using the information of the phylogeny of the systems, we added sequence of systems that was annotated as generic but that clustered in a group of experimentally validated systems. We aligned all the representatives using MAFFT v7.273 (--auto, automatic selection of the parameters depending of the size of the alignment, default values for the other parameters) [100]. With SEAVIEW [105], the poorly aligned regions at the extremities were manually trimmed in the alignment. The trimmed alignment was used to build the HMM profile using hmmbuild (default parameters) from HMMER package v3.1b2 [106].

### Phylogenetic inference

Phylogenetic analyses based on protein sequences involved an initial alignment of the sequences using MAFFT v7.273 (“linsi” algorithm) [100]. Multiple alignments were analysed using Noisy v1.5.12 (default parameters) [107] to select the informative sites. We inferred maximum likelihood trees from the curated alignments, or their concatenates, using IQ-TREE v 1.6.7.2 [51] (options -allnni, -nstop 1000, -nm 100000). We evaluated the node supports using the options -bb 1000 for ultra-fast bootstraps and -alrt 1000 for SH-aLRT [108]. The best evolutionary model was selected with ModelFinder (option -MF, BIC criterion) [109]. We used the option -wbtl to conserve all optimal trees and their branch lengths.

The phylogenetic trees of 16S rRNA sequences were built from a dataset including one sequence per genome of 5776 genomes. The 16S sequences were retrieved from genome sequences using RNammer v1.2 [110] (options -S set to bac and the – m to ssu). We aligned separately archaeal and bacterial 16S rRNA using the secondary structure models with SSU_Align v0.1.1 (http://eddylab.org/software/ssu-align/, default options). Poorly aligned positions were masked with ssu-mask. The alignment was trimmed with trimAl v1.4rev15 [111] (-noallgaps, that allow to remove from the alignment regions that are only composed of gaps). The maximum likelihood trees were inferred using IQ-TREE v1.6.7.2 [51] (using the best-selected model SYM+R6 for the archaeal tree and SYM+R10 for the bacterial tree, -bb 10000 ultrafast bootstrap [108], -wbtl to conserve all optimal trees and their branch lengths).

### Reference systems dataset

The dataset with all the systems identified in the genomes is too large to make phylogenetic inferences. It also contains many very closely related systems that may provide little additional information to infer the deeper nodes of the tree. Hence, we developed a method to remove redundancy in the dataset while maximising its genetic diversity. The method prioritizes the inclusion of systems that were experimentally validated to facilitate the analysis of the results. The method consists of several sequential steps (Figure S15).
1. We inferred the maximum likelihood tree for each key components’ family as mentioned above and extracted the matrices of patristic distances (using the R function “cophenetic_phylo” of the package ape) between all leaves of the trees. This resulted in a set of distance matrices between systems.
2. When there were multiple copies of family of clade-specific paralogs the system was represented multiple times in the phylogeny and in the distance matrix. To solve the problem and to have only one distance between two systems, we chose the minimal distance between the two systems between paralogs.
3. Each core protein family has a different rate of evolution. To compare them, we normalized each matrix by the sum of all the branch lengths in the tree of the family. We then built a matrix that is the average of all normalized matrices. This average matrix was used to infer a tree with bioNJ [112]. The tree was rooted at midpoint.
4. We used the bioNJ tree to define monophyletic groups of similar systems. We iteratively used the R function “cutree” from the stats package by gradually decreasing or increasing the heights where the tree should be “cut” until we obtained between 200 and 300 groups.
5. At this stage in the method, we had obtained a set of monophyletic groups of closely related systems. To pick the representative system of each group, we had the following order of priorities: i) inclusion of systems validated experimentally, ii) inclusion of the systems with fewest paralogs. In some rare cases, a given type of systems (*e.g.*, T2SS) had less than 20 instances after this procedure. In this case, and to increase the statistical power of the analyses, we modified the height of the “cutree” function for the specific subtree of the systems lacking representative to obtain a minimum of 20 systems for each group if possible. The systems selected to make the phylogeny are named “representative systems”.
6. We removed some complex systems from the reference ones (6/39 Archaeal-T4P and 10/101 generic) because they had two paralogs of all the genes or were punctual generic systems with components from different types (for example: T2SS_gspE and T4P_pilB).

### De-replicated dataset

To reduce the number of paralogs in each system, we used the following method (Fig. S16).
1. We inferred the maximum likelihood tree for each key components’ family of the representative dataset as mentioned above and extracted the matrices of patristic distances (using the R function “cophenetic_phylo” of the package ape) between all leaves of the trees. This resulted in a set of distance matrices between proteins.
2. For each system with more than one copy per gene, we found the nearest system based on patristic distances extracted from the ATPase or the integral membrane platform tree (depending on the number of copies of the ATPase), which had only one copy of this gene.
3. We use this nearest system to choose the copy of the duplicate gene with the smallest distance to its homolog in this nearest system.
4. In the end, each system is represented by a single instance of each of the core proteins and we called this set of selected sequences and systems the “dereplicated dataset”.

### Concatenate trees and ML topology tests

The Tad pilus and T4P both show cases of system-wide duplications: some of their gene families harbour paralogs issued from ancestral duplications, as supported from individual gene trees analyses. That is the case of the IM-platform for the Tad pilus (TadB and TadC are paralogs derived from an ancestral duplication) and of the ATPase for the T4P (PilB and PilT/PilU are paralogs). Candidate systems’ trees were generated based on the concatenation of all possible combinations of putative sets of orthologs, i.e., each paralog was picked one after the other to represent their system in phylogenetic analyses. For instance, for the IM-platform, a first set of orthologs would consist of TadB sequences for Tad systems together with IM-platform sequences from all other systems, and another would consist of TadC sequences for Tad systems together with IM-platform sequences from all other systems. For the ATPase, we decided to only focus on the functional orthologous gene, the PilB sequences for T4aP systems.

Therefore, there were two possible combinations of the mandatory genes to generate concatenates of the ATPase, IM-platform. In total, we generated two concatenates, and used IQ-Tree to compute maximum-likelihood phylogenetic trees, using partition models (option -spp, the location of the genes in the concatenation define the partitions, the model for each partition corresponds to the model found previously for the individual analyses).

In order to assess the congruence between the concatenate trees and the individual protein trees, a maximum-likelihood topology test (AU for “Approximately Unbiased”,[59] was performed using IQ-Tree. Each protein alignment was used as an input to assess the congruence of its ML tree with those of the ten concatenate trees. The parameters of the model were estimated on the initial parsimony tree (option -n 0). A correction for multiple tests was applied to the p-values (sequential Bonferroni per batch of ten concatenate trees).

### Analysis of the neighbourhood of the systems

We searched for genes systematically associated with a given type of systems by analysing the neighbourhood of each system. For each locus of a system we identified its first gene (position X_First_) and last gene (position X_End_). We then took all the genes in a neighbourhood of 10 (i.e. between X_First-10_ and X_End+10_). When a system was encoded in multiple loci in the genome, each locus was analysed in the same way. We then clustered all these proteins by sequence similarity using BLASTP v. 2.5.0+ (default settings, e-value < 10^−6^) [103] and Silix v1.2.10-p1 (minimal percentage of identity to accept blast hits for building families at 0.5) [104]. We kept the clusters if they had proteins represented in systems of different leaves in the tree. The proteins of each cluster were aligned with MAFFT and used to build HMM protein profiles as described above.

To annotate the protein clusters we used two methods. First, we searched for similarities of their HMM profiles with the profiles used to identify the TFF-SF components using HHsearch (v3.0.3, p-value < 10^−5^). Second, we searched for homologies between the remaining clusters and the profiles of the PFAM database (v31.0, same method).

To test if a given cluster is significantly positively associated with a given type of system, we made the following analysis. We counted the occurrences of the elements of the cluster associated with a given type and made a contingency table where the columns are the type versus all other types and the lines are presence or absence of an element from the cluster. To test statistical significance, we used a Fisher’s exact test on the contingency table. Since this implicates many statistical tests – one test per type per cluster and this for many clusters and several types – we adjusted the p-values for multiple tests using the Bonferroni correction. We kept the association between a given cluster and a given type of system if the number of elements in the cluster neighbouring systems of that type was higher than expected by chance and if the corrected p-value < .0.05. The resulting matrix of presence/absence for genes positively associated with the systems can be found in the Table S5.

### Inferring transfers of systems

We took the phylogenetic tree of all systems and picked the sub-trees of each type of system. For each of these trees, we pruned the 16S rRNA tree such that it only includes species present in the system tree. We used ALE v0.4 (default parameters) a reconciliation program which introduces events of duplications, transfers and losses (DTL) in a gene tree, and amalgamate most frequent subtrees in a sample distribution of the gene tree to improve it, and make it congruent with the species (reference) tree in a maximum-likelihood framework [113]. Using ALE we computed a number of DTL events introduced in each system’s trees given 16S rRNA tree as a reference. We then collected the number of transfers for each type of system and divided this number by the number of branches in the sub-trees of each type of system to have the proportion of transfers.

### Analysis of genetic organisation

We identified all pairs of contiguous components of the systems (for a gene X_p_ in the cluster we look at the genes X_p-1_ and X_p+1_). We constructed an adjacency matrix using this information, and we used it to construct a graph of the genetic organisation of the systems. We normalized the association between two genes to represent two different types of information:

1. To know how frequently two genes are contiguous, we divided the number of contiguous occurrences by the number of occurrences of the rarest of the two genes. This corresponds to edges’ width in Fig 5.
2. To know how many times the contiguity is found in the system, we divided the number of times the contiguity is observed by the number of systems detected. This corresponds to edges’ colour in Fig 5.

## Data and software availability

The data produced by these analyses are available at: https://gitlab.pasteur.fr/rdenise/diversification-of-tff-sf-data, accessed March 2019.

## Author contributions

Conceptualization: RD, SA, EPCR

Data curation: RD

Funding acquisition: RD, EPCR

Investigation: RD, SA, EPCR

Methodology: RD, SA, EPCR

Software: RD, SA

Supervision: SA, EPCR

Writing – original draft: RD, SA, EPCR

Writing – review & editing: RD, SA, EPCR

## Acknowledgements

We are much indebted to Olivera Francetic for comments, encouragement and guidance during the development of this study. We thank Simonetta Gribaldo for comments and suggestions on a previous version of the manuscript, Bertrand Néron for continuous support of MacSyFinder. We also thank Panagiotis Adam for providing a comprehensive phylogenetic tree of the archaeal and bacterial domains to be used as support for Figs. 4, S2 and S11.

## Supplemental Fig legends

**Fig S1. Representation of the initial and final models of the systems.** Homologous genes are indicated by a same colour. Mandatory genes are indicated with a full outline, accessory genes are indicated with a dash outline and forbidden genes are indicated with a red cross. The exchangeable genes are indicated by an arrow. The loner genes are indicated by a star below the gene. For the Archaeal-T4P the aCXXX name indicates that all the homologous arCOGs for this function (they are set as exchangeable). The empty box in the genetic model indicates that the genes are exchangeable with all the homologous genes of the other models.

**Fig S2. Taxonomic distribution of the systems in Bacteria and Archaea with the initial models.** Cells indicate the number of genomes with at least one detected system. The cell’s colour gradient represents the proportion of genomes with at least one system in the clade. The bar plot shows the total number of detected systems. The bars are separated in two categories: Alpha-, Beta-, Gamma-proteobacteria versus the other clades. The cladogram symbolizes approximated relationships between the bacterial and archaeal taxa analysed in this study.

**Fig S3. Rooted phylogeny of the ATPase.** The colour of the label of the leaves indicates the taxonomic group of the species. The different coloured strips indicate the classification of the systems with the MacSyFinder annotation (with the initial model and with the final one) and the annotation of the systems in the literature. The systems known to be implicated in natural transformation are indicated in dark purple. Known sub-types of Archaeal-T4P are indicate by a text in red. The annotation of the domains of the proteins using are also added. The tree was built using IQ-Tree, 10000 replicates of UF-Boot, model LG+10.

**Fig S4. Rooted phylogeny of the ATPase with FtsK as external group.** The colour of the label of the leaves indicates the taxonomic group of the species. The different coloured strips indicate the classification of the systems with the MacSyFinder annotation (with the initial model and with the final one) and the annotation of the systems in the literature. The systems known to be implicated in natural transformation are indicated in dark purple. Known sub-types of Archaeal-T4P are indicate by a text in red. The annotation of the domains of the proteins using are also added. The tree was built using IQ-Tree, 10000 replicates of UF-Boot, model LG+R10.

**Fig S5. Rooted phylogeny of the ATPase with FtsK and virB4 as external group.** The colour of the label of the leaves indicates the taxonomic group of the species. The different coloured strips indicate the classification of the systems with the MacSyFinder annotation (with the initial model and with the final one) and the annotation of the systems in the literature. The systems known to be implicated in natural transformation are indicated in dark purple. Known sub-types of Archaeal-T4P are indicate by a text in red. The annotation of the domains of the proteins using are also added. The tree was built using IQ-Tree, 10000 replicates of UF-Boot, model LG+R9.

**Fig S6. Rooted phylogeny of the integral membrane platform.** The colour of the label of the leaves indicates the taxonomic group of the species. The different coloured strips indicate the classification of the systems with the MacSyFinder annotation (with the initial model and with the final one) and the annotation of the systems in the literature. The systems known to be implicated in natural transformation are indicated in dark purple. Known sub-types of Archaeal-T4P are indicate by a text in red. The annotation of the domains of the proteins using are also added. The tree was built using IQ-Tree, 10000 replicates of UF-Boot, model LG+F+R8.

**Fig S7. Rooted phylogeny of the major pilin.** The colour of the label of the leaves indicates the taxonomic group of the species. The different coloured strips indicate the classification of the systems with the MacSyFinder annotation (with the initial model and with the final one) and the annotation of the systems in the literature. The systems known to be implicated in natural transformation are indicated in dark purple. Known sub-types of Archaeal-T4P are indicate by a text in red. The annotation of the domains of the proteins using are also added. The tree was built using IQ-Tree, 10000 replicates of UF-Boot, model LG+F+R7.

**Fig S8. Unrooted phylogeny of the prepilin peptidase.** The colour of the label of the leaves indicates the taxonomic group of the species. The different coloured strips indicate the classification of the systems with the MacSyFinder annotation (with the initial model and with the final one) and the annotation of the systems in the literature. The systems known to be implicated in natural transformation are indicated in dark purple. Known sub-types of Archaeal-T4P are indicate by a text in red. The annotation of the domains of the proteins using are also added. The tree was built using IQ-Tree, 10000 replicates of UF-Boot, model VT+F+R6.

**Fig S9. Unrooted phylogeny of the secretin.** The colour of the label of the leaves indicates the taxonomic group of the species. The different coloured strips indicate the classification of the systems with the MacSyFinder annotation (with the initial model and with the final one) and the annotation of the systems in the literature. The systems known to be implicated in natural transformation are indicated in dark purple. Known sub-types of Archaeal-T4P are indicate by a text in red. The annotation of the domains of the proteins using are also added. The tree was built using IQ-Tree, 10000 replicates of UF-Boot, model LG+F+R8.

**Fig S10. Rooted phylogeny of the TFF-SF.** The was built with the concatenate of the IM platform (using TadB) and the AAA+ ATPase (using PilB). The colour of the label of the leaves indicates the taxonomic group of the species. The different coloured strips indicate the classification of the systems with the MacSyFinder annotation (with the initial model and with the final one) and the annotation of the systems in the literature. The systems known to be implicated in natural transformation are indicated in dark purple. Known sub-types of Archaeal-T4P are indicate by a text in red. The tree was built using IQ-Tree, 10000 replicates of UF-Boot, with a partition model.

**Fig S11. Taxonomic distribution of the systems in Bacteria and Archaea using the phylogenetic clustering to annotate generic systems.** Cells indicate the number of genomes with at least one detected system. The cell’s colour gradient represents the proportion of genomes with at least one system in the clade. The bar plot shows the total number of detected systems. The bars are separated in two categories: Alpha-, Beta-, Gamma-proteobacteria versus the other clades. The cladogram symbolizes approximated relationships between the bacterial and archaeal taxa analysed in this study.

**Fig S12. Genetic organization of the Archaeal-T4P in genomes.** The edge width represents the number of times the two genes are contiguous divided by the number of times the rarest gene is present in the system. The colour of the edge represents the number of times the two genes are contiguous in the system divided by the number of systems.

**Fig S13. Genetic organization of the Archaellum in genomes.** The edge width represents the number of times the two genes are contiguous divided by the number of times the rarest gene is present in the system. The colour of the edge represents the number of times the two genes are contiguous in the system divided by the number of systems.

**Fig S14. 16S tree used to infer horizontal transfers.** The colour of the leaves represents the phyla of the bacteria. The tree was built using IQ-Tree, 1000 replicates of UF-Boot, model SYM+R10.

**Fig S15. Schema of the workflow used to choose the representative systems.**

**Fig S16. Schema of the workflow used to choose the species-specific paralogs.**

## Supplemental Table legends

Table S1. List of all the profiles of the TFF-SF used in the analysis.

Table S2. Experimentally validated systems used in the analysis.

Table S3. Description of all the genes and concatenate trees inferred in this study.

Table S4. All trees inferred in this study in newick format.

Table S5. Matrix of presence/absence of neighbouring genes positively associated with the systems (Family of genes are in columns and systems are in rows).

Table S6. All the systems detected by Macsyfinder with the search using the final models.

Table S7. Tree topology tests (AU) using IQ-TREE between concatenated trees and the genes that compose the concatenate.

