## Supplementary figures and images for "Diversification of the type IV filament super-family into machines for adhesion, secretion, DNA transformation and motility"

### Supplemental Figure 1

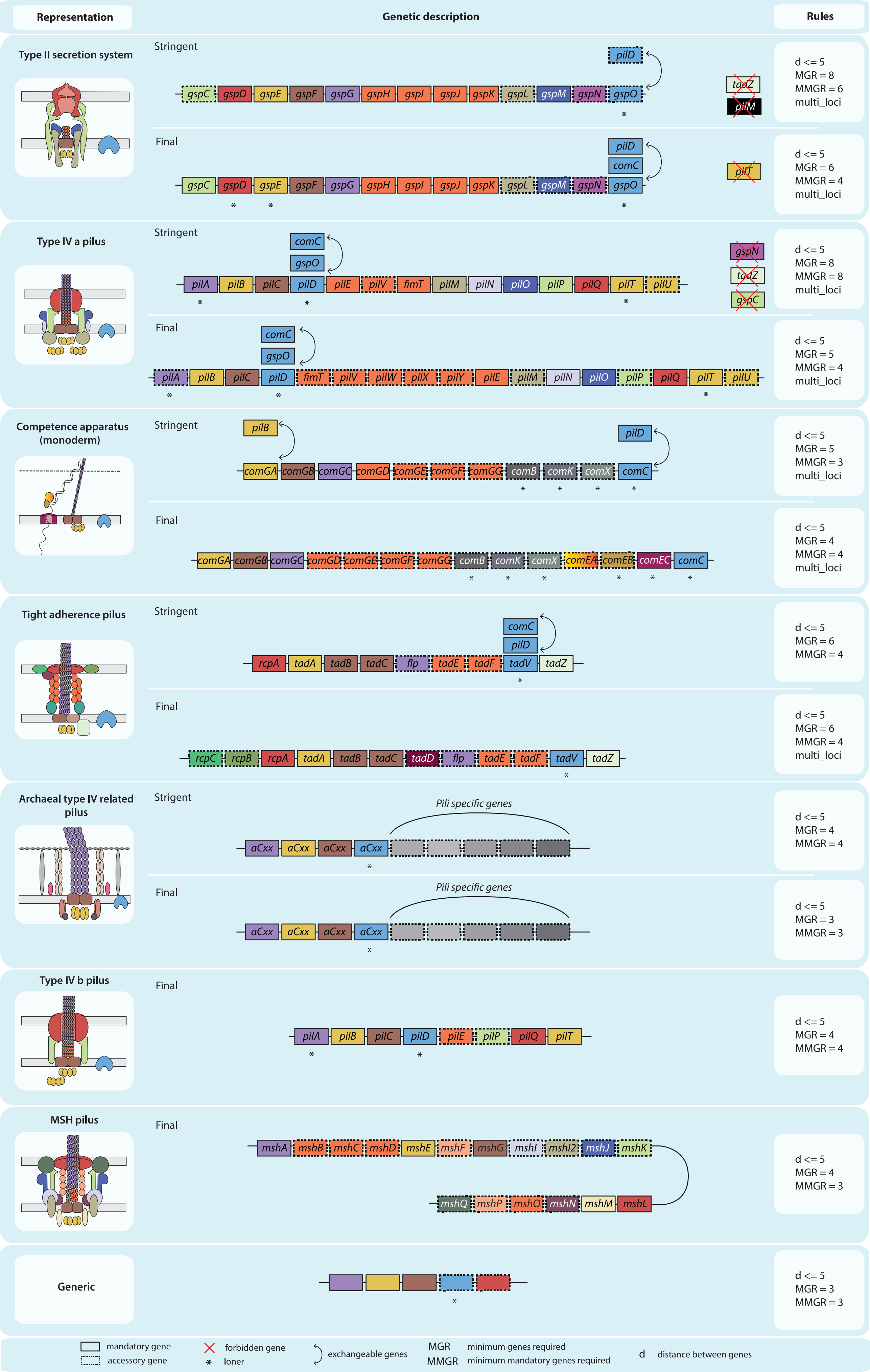

### Supplemental Figure 3

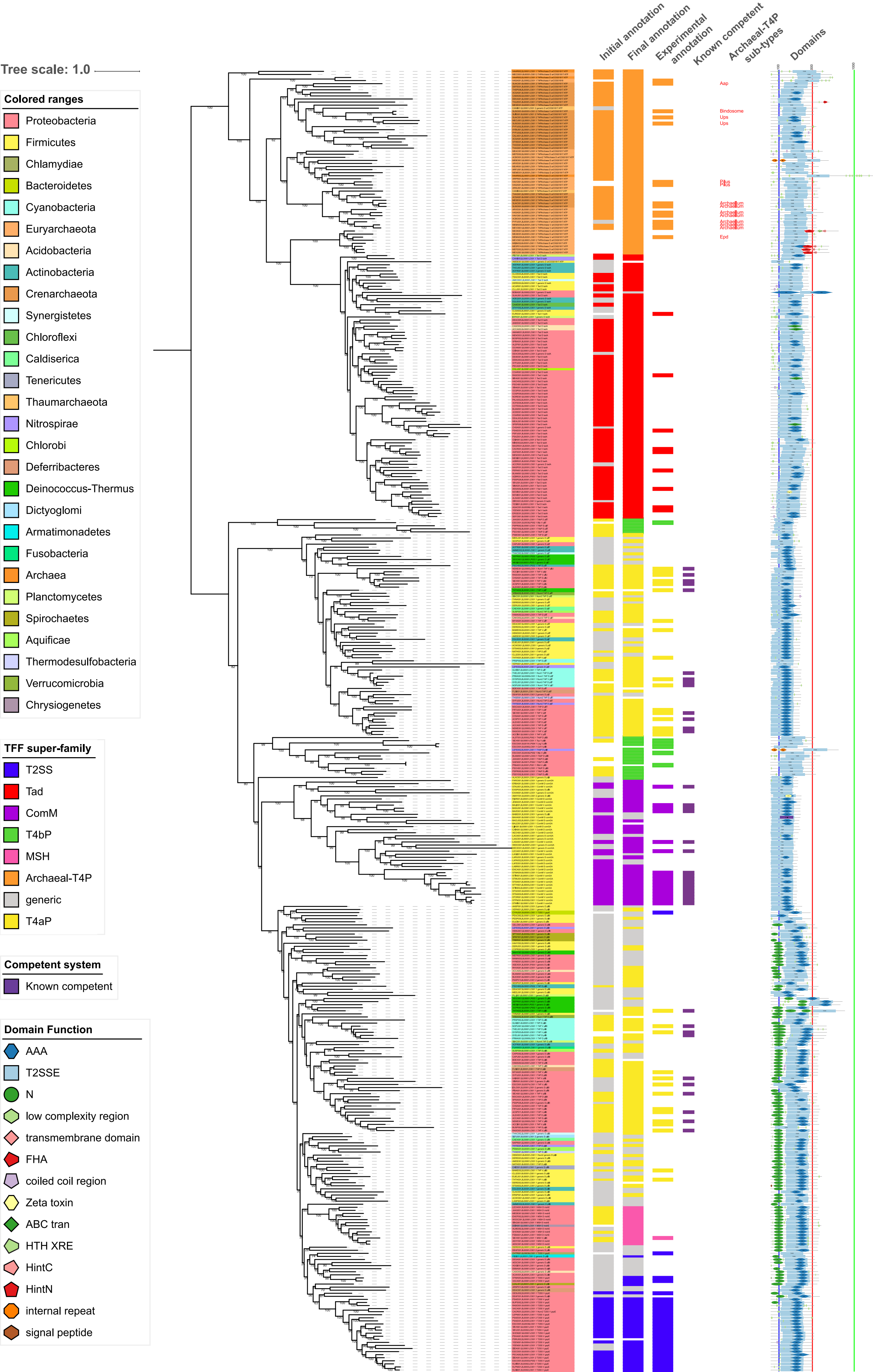

### Supplemental Figure 4

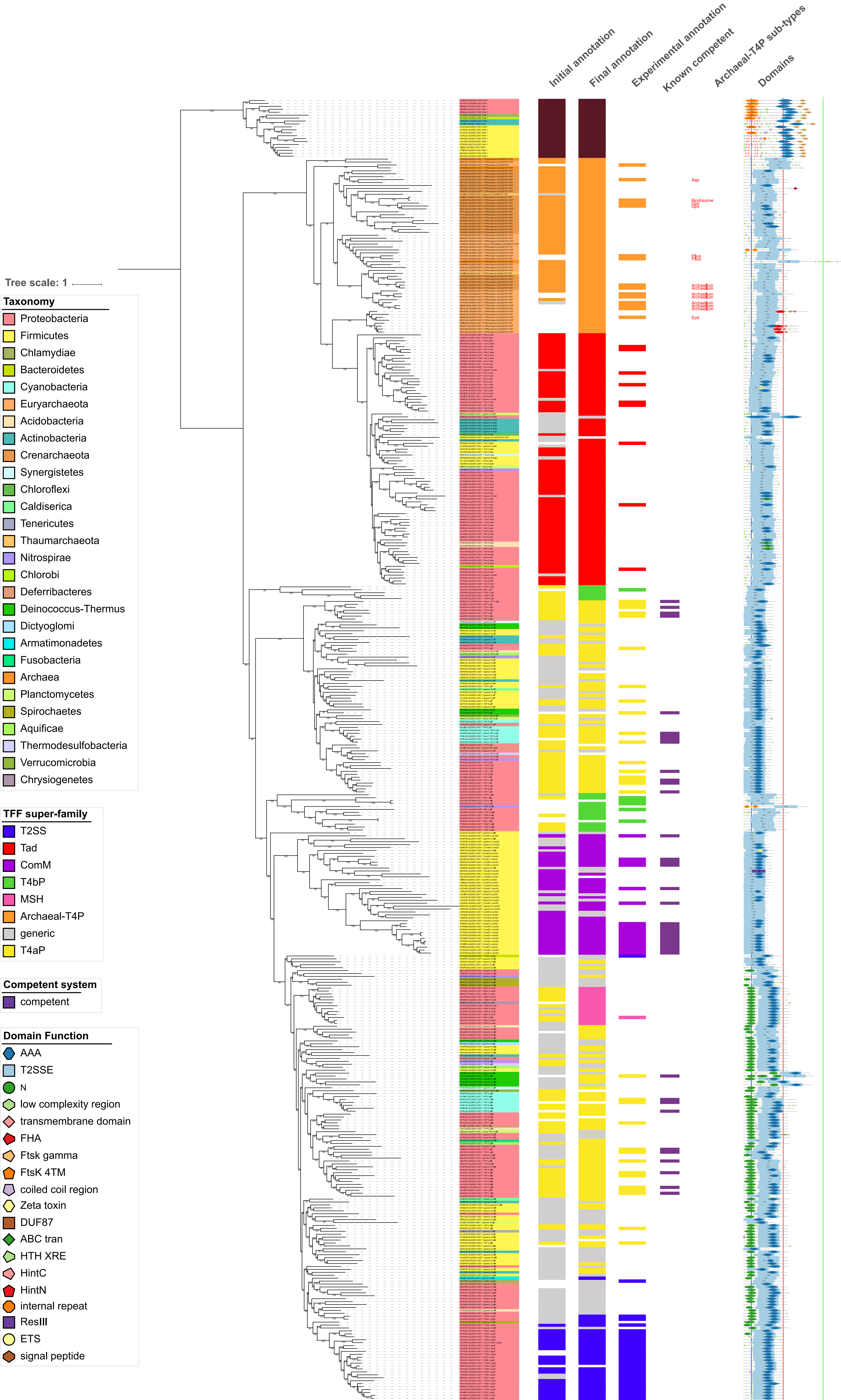

### Supplemental Figure 5

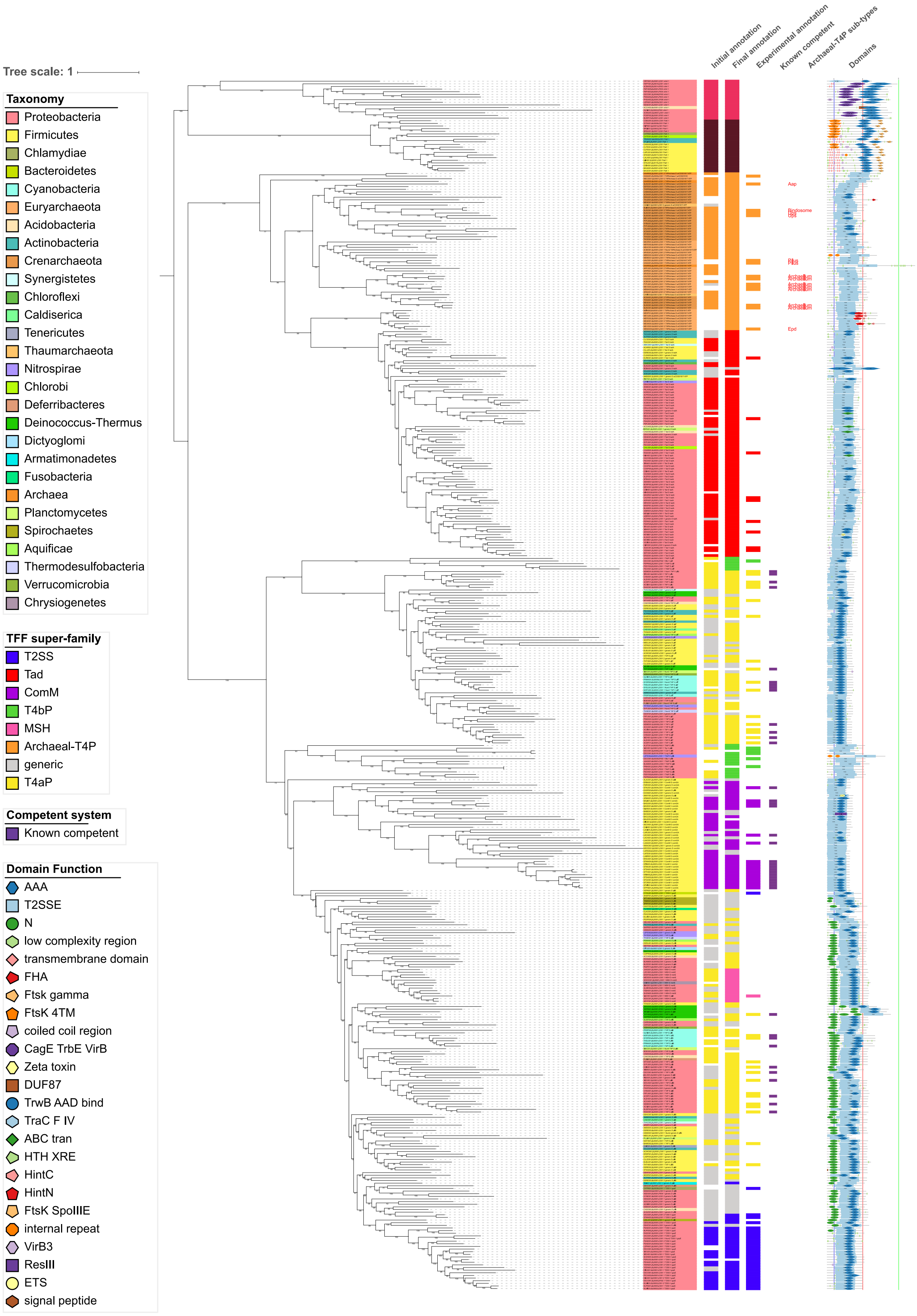

### Supplemental Figure 6

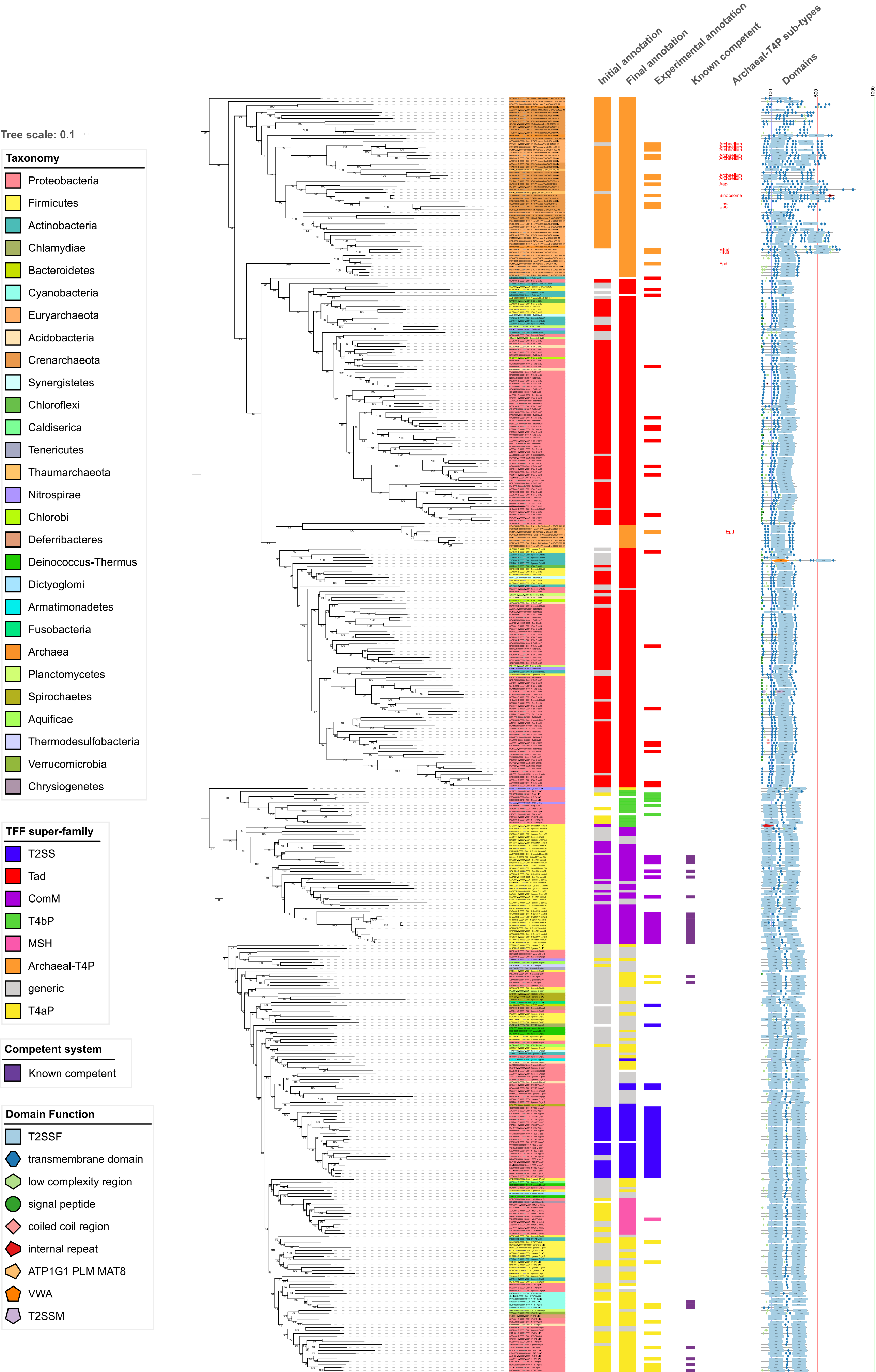

### Supplemental Figure 7

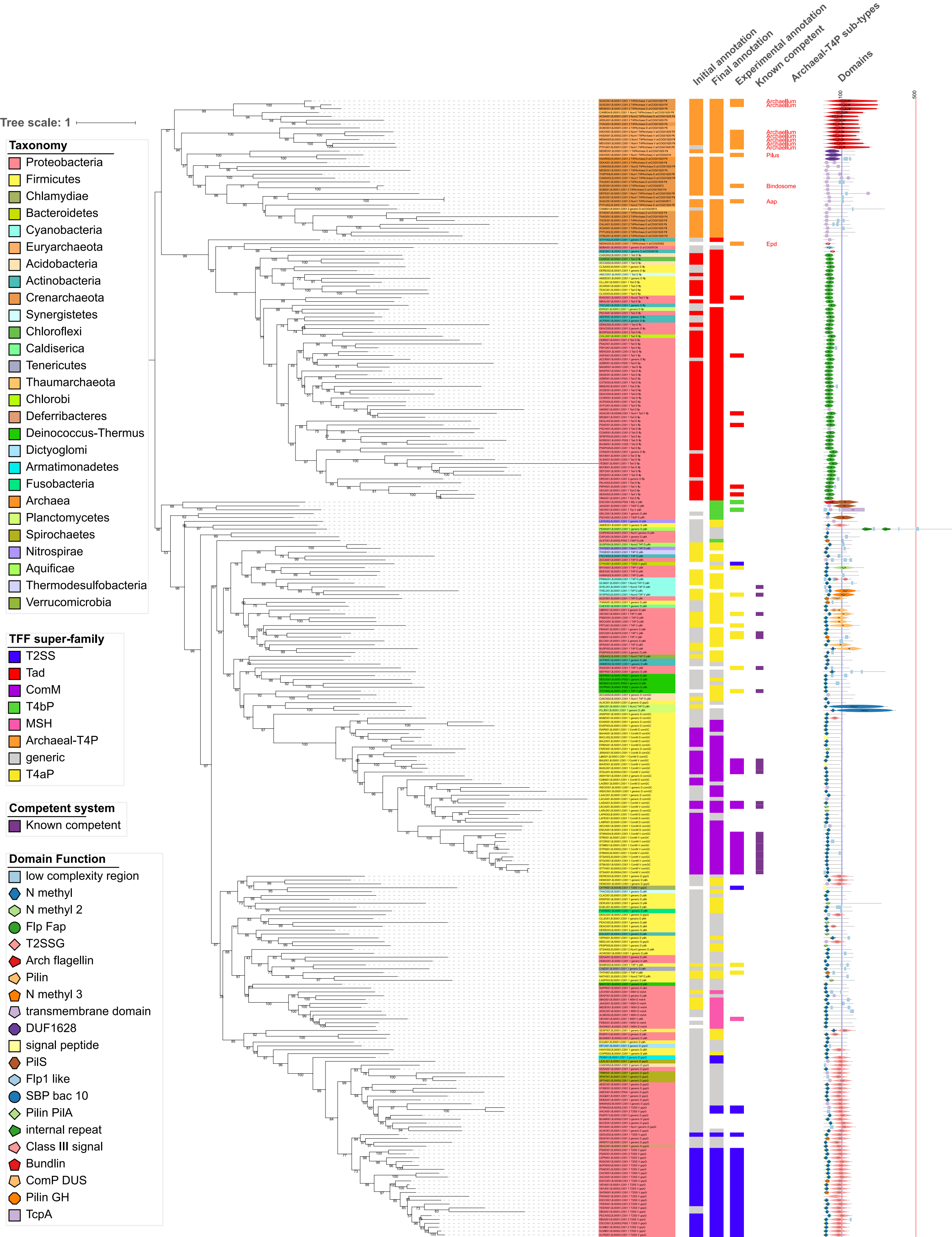

### Supplemental Figure 8

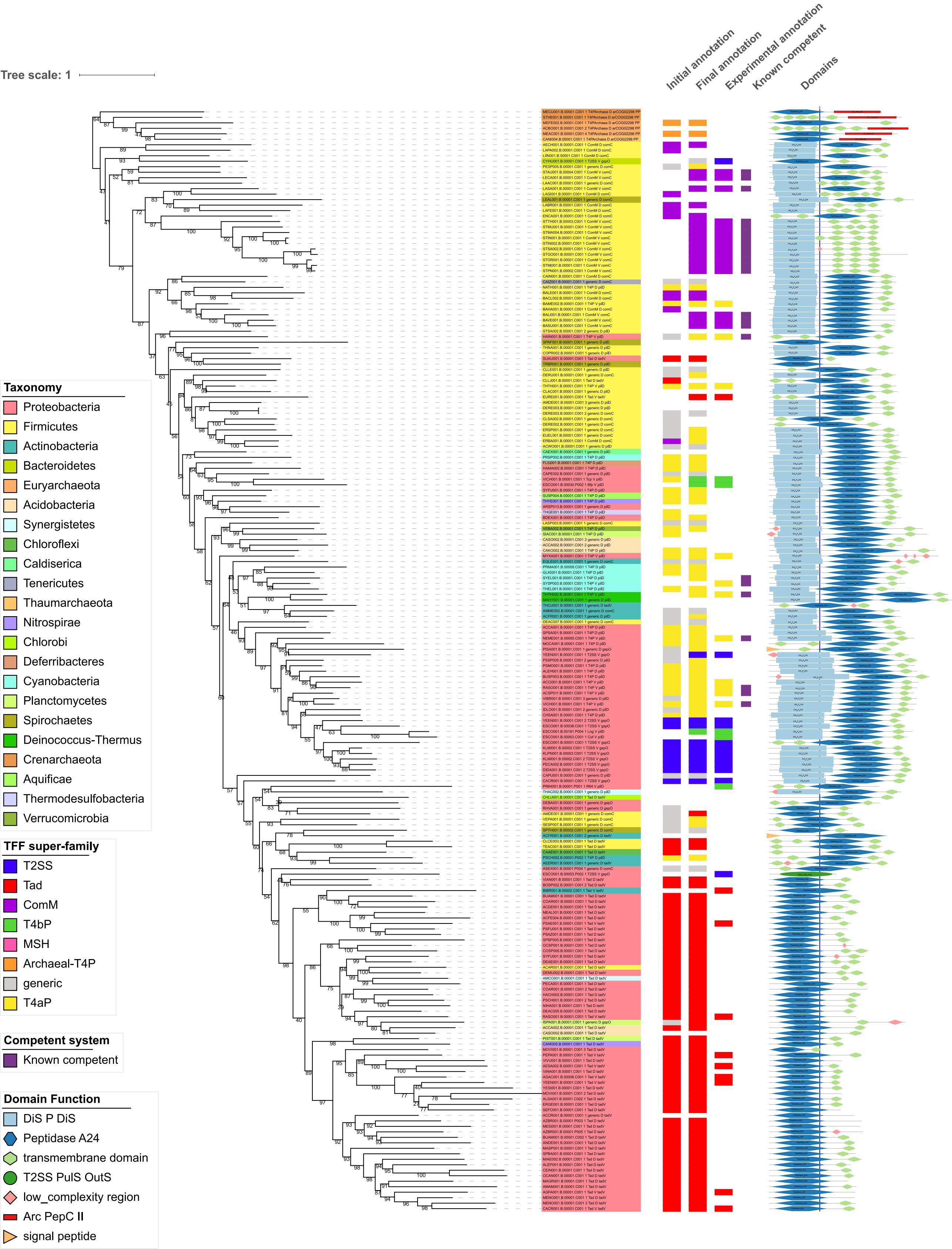

### Supplemental Figure 9

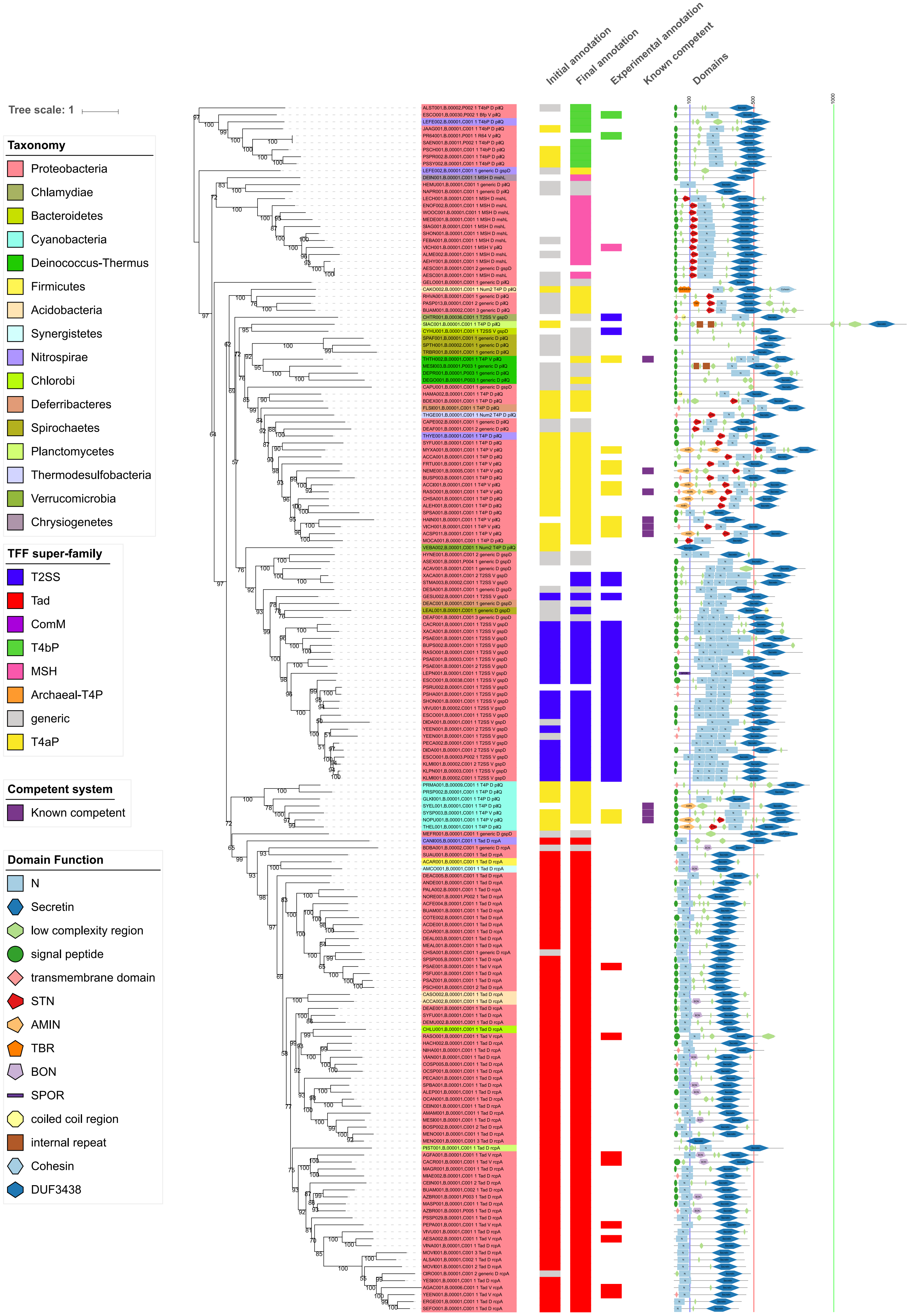

### Supplemental Figure 10

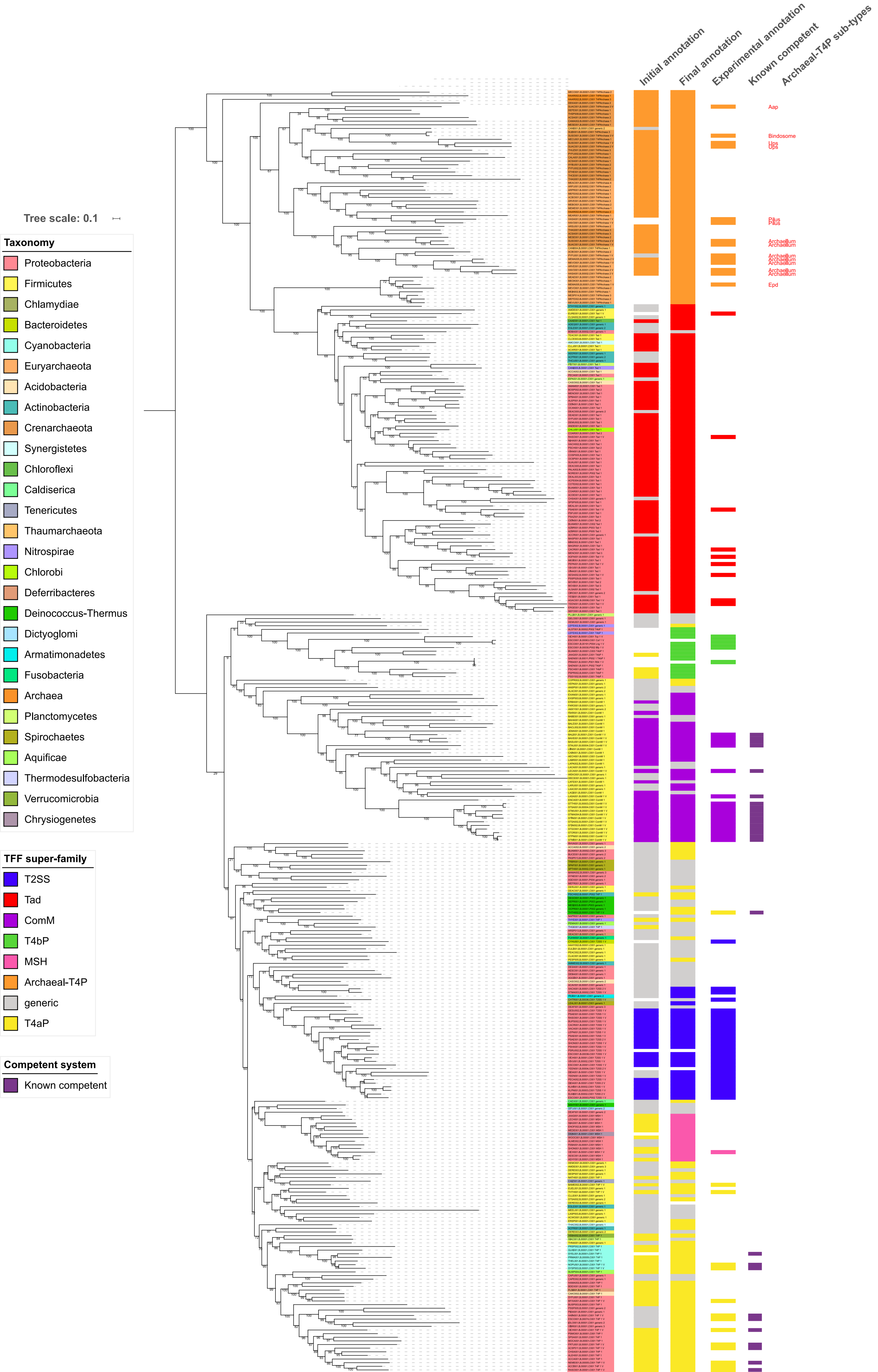

### Supplemental Figure 12

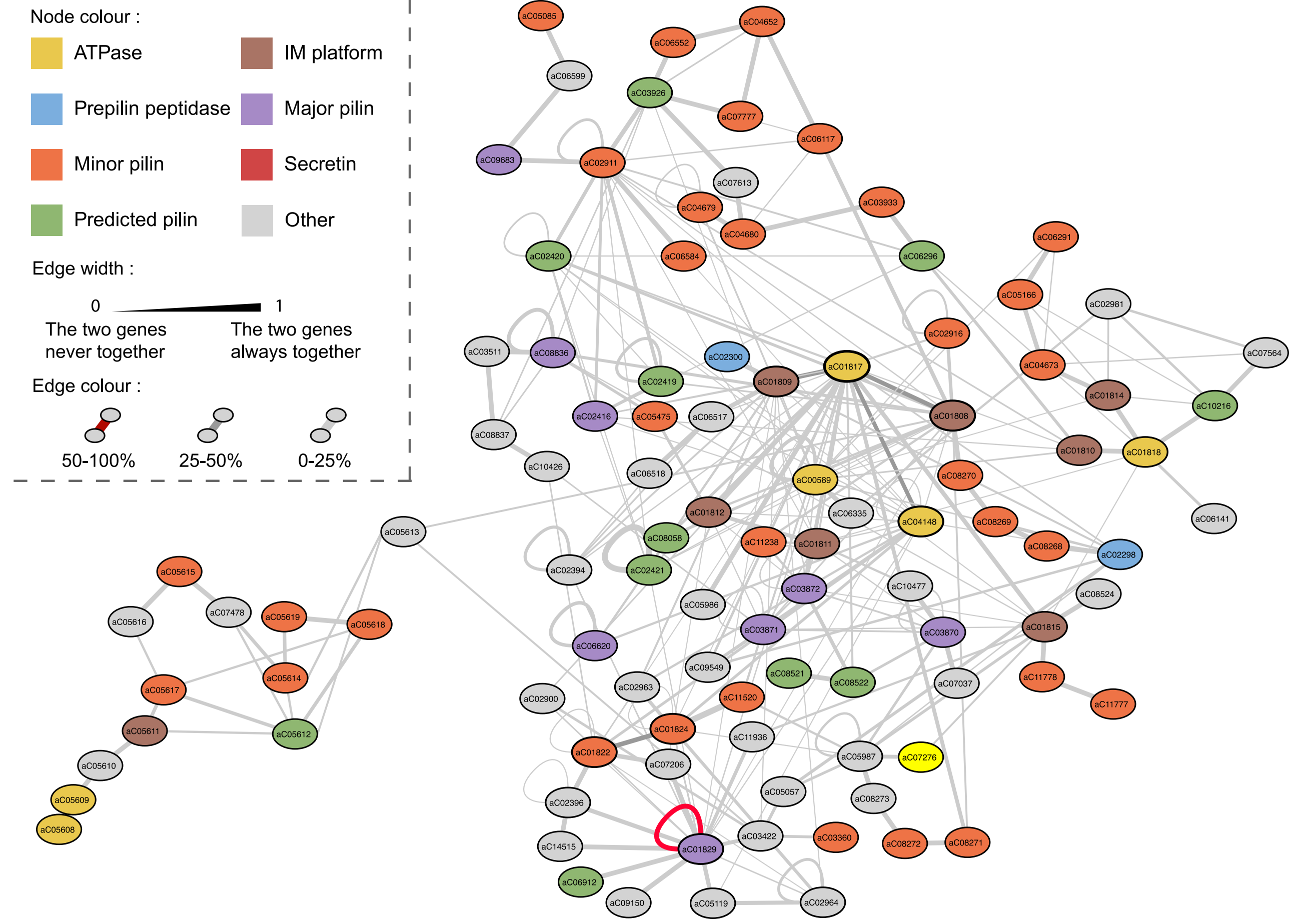

### Supplemental Figure 13

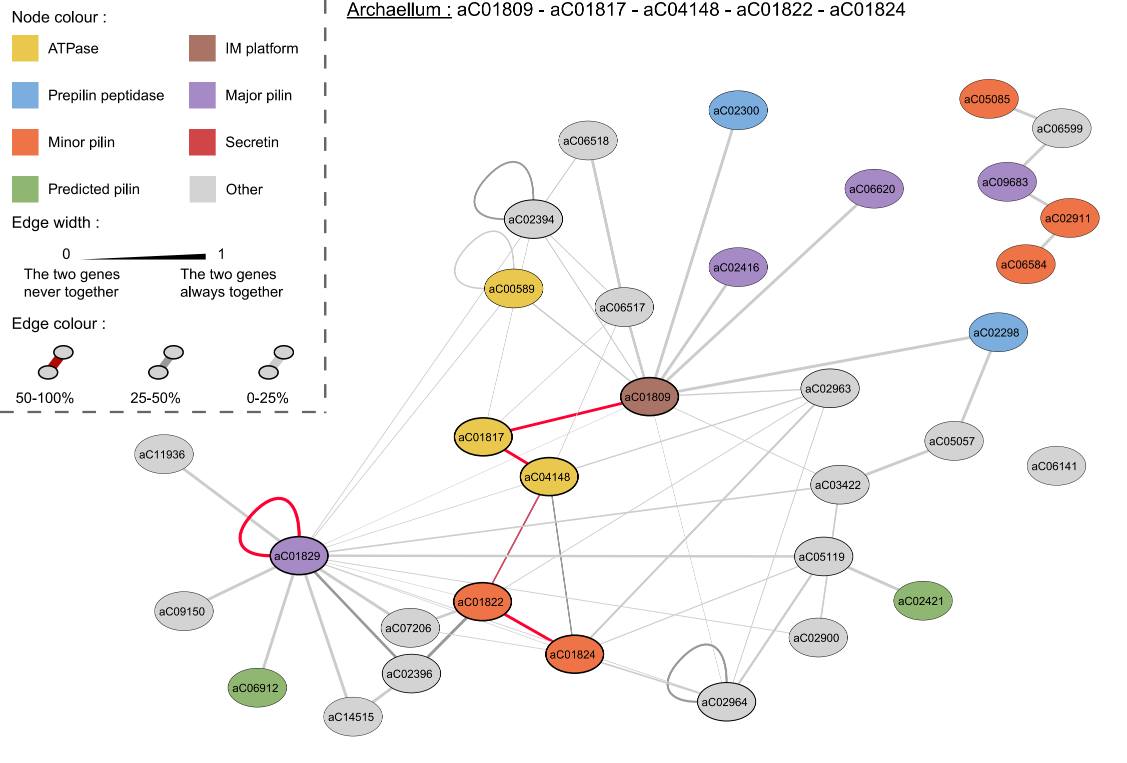

### Supplemental Figure 14

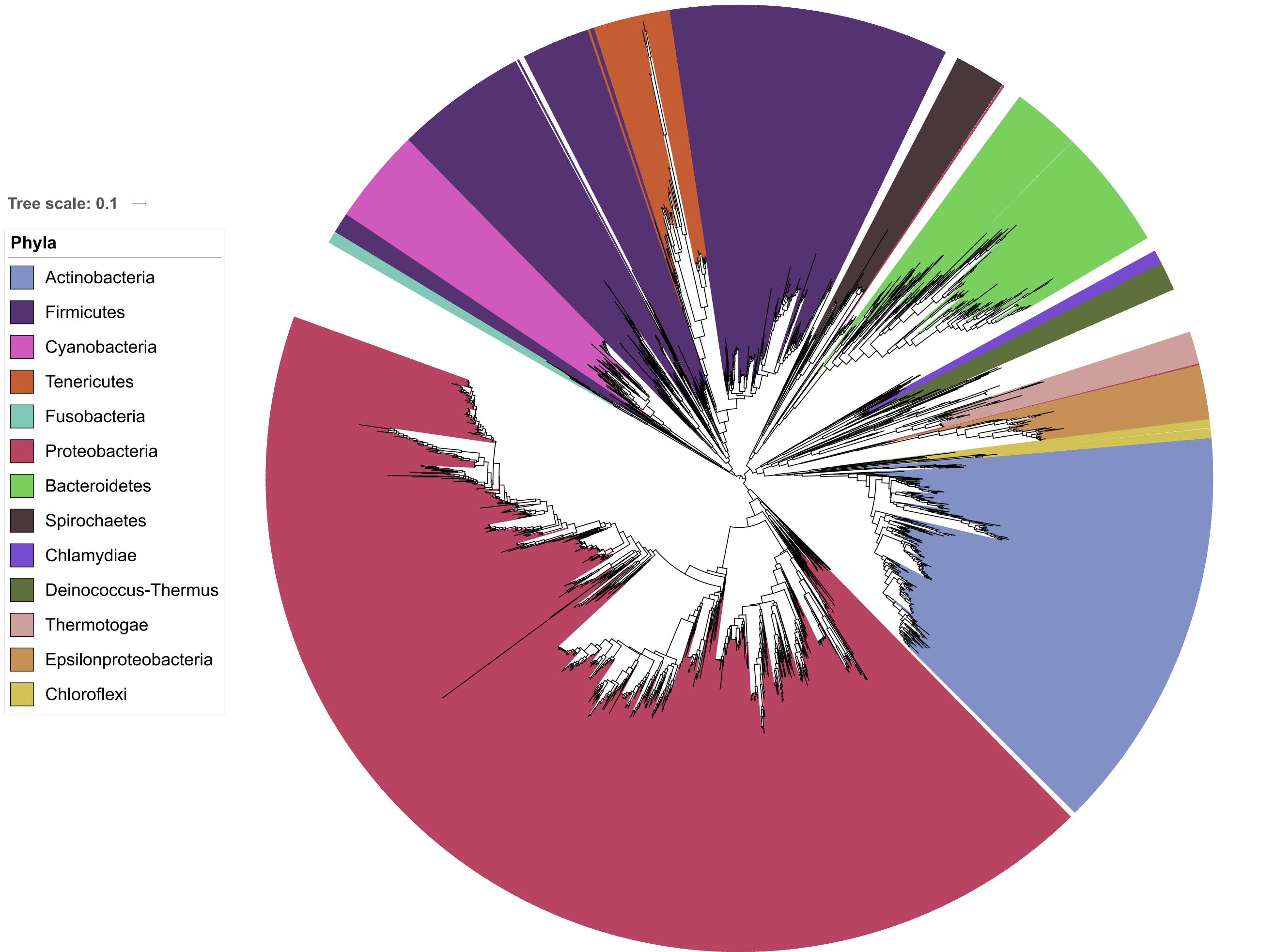
