## Supplemental Figure 2 for "Diversification of the type IV filament super-family into machines for adhesion, secretion, DNA transformation and motility"

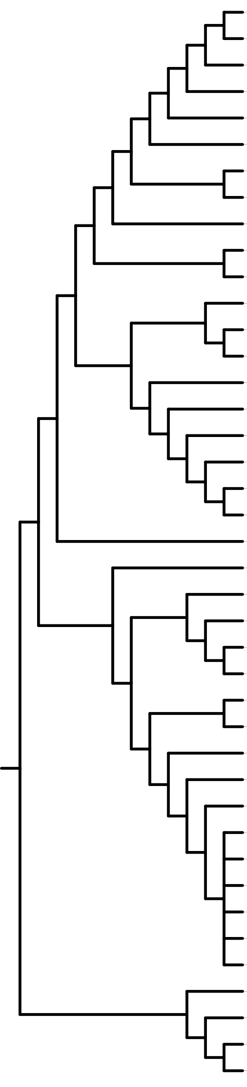

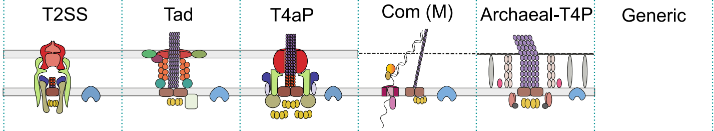


| Taxon | Number of genomes | T2SS | Tad | T4aP | Com (M) | Archaeal-T4P | Generic |
|---|---|---|---|---|---|---|---|
| Gammaproteobacteria | 1676 | 901 | 367 | 720 | 0 | 0 | 621 |
| Betaproteobacteria | 500 | 259 | 314 | 206 | 0 | 0 | 48 |
| Acidithiobacillia | 5 | 0 | 2 | 5 | 0 | 0 | 0 |
| Alphaproteobacteria | 469 | 39 | 219 | 0 | 0 | 0 | 33 |
| Deltaproteobacteria | 75 | 11 | 25 | 47 | 0 | 0 | 33 |
| Nitrospirae | 8 | 0 | 2 | 4 | 0 | 0 | 7 |
| Deferribacteres | 4 | 0 | 0 | 3 | 0 | 0 | 2 |
| Chrysiogenetes | 1 | 1 | 0 | 0 | 0 | 0 | 0 |
| Acidobacteria | 8 | 0 | 4 | 2 | 0 | 0 | 1 |
| Epsilonproteobacteria | 264 | 0 | 1 | 0 | 0 | 0 | 8 |
| Aquificae | 16 | 0 | 0 | 2 | 0 | 0 | 5 |
| Planctomycetes | 8 | 0 | 2 | 1 | 0 | 0 | 4 |
| Chlamydiae | 74 | 0 | 0 | 0 | 0 | 0 | 22 |
| Verrucomicrobia | 12 | 0 | 0 | 1 | 0 | 0 | 0 |
| Elusimicrobia | 3 | 0 | 0 | 0 | 0 | 0 | 1 |
| Gemmatimonadetes | 1 | 0 | 0 | 0 | 0 | 0 | 0 |
| Fibrobacteres | 2 | 0 | 0 | 2 | 0 | 0 | 0 |
| Ignavibacteria | 2 | 0 | 0 | 0 | 0 | 0 | 0 |
| Chlorobi | 13 | 0 | 6 | 0 | 0 | 0 | 0 |
| Bacteroidetes | 194 | 0 | 0 | 0 | 0 | 0 | 4 |
| Spirochaetes | 66 | 0 | 0 | 0 | 0 | 0 | 19 |
| Fusobacteria | 20 | 0 | 0 | 1 | 0 | 0 | 12 |
| Synergistetes | 5 | 0 | 1 | 0 | 0 | 0 | 2 |
| Deinococcus-Thermus | 26 | 0 | 0 | 1 | 0 | 0 | 9 |
| Thermotogae | 32 | 0 | 0 | 0 | 0 | 0 | 0 |
| Caldiserica | 1 | 0 | 0 | 0 | 0 | 0 | 1 |
| Dictyoglomi | 2 | 0 | 0 | 2 | 0 | 0 | 0 |
| Cyanobacteria | 94 | 0 | 0 | 55 | 0 | 0 | 0 |
| Chloroflexi | 25 | 0 | 1 | 0 | 0 | 0 | 0 |
| Actinobacteria | 608 | 0 | 15 | 3 | 0 | 0 | 44 |
| Armatimonadetes | 1 | 0 | 0 | 1 | 0 | 0 | 0 |
| Bacilli | 1018 | 0 | 4 | 34 | 872 | 0 | 82 |
| Clostridia | 158 | 0 | 14 | 45 | 0 | 0 | 74 |
| Erysipelotrichia | 8 | 0 | 3 | 0 | 1 | 0 | 3 |
| Negativicutes | 12 | 0 | 1 | 2 | 0 | 0 | 3 |
| Tissierellia | 4 | 0 | 0 | 0 | 0 | 0 | 1 |
| Tenericutes | 141 | 0 | 0 | 0 | 0 | 0 | 1 |
| Euryarchaeota | 155 | 0 | 0 | 0 | 0 | 101 | 23 |
| Korarchaeota | 1 | 0 | 0 | 0 | 0 | 0 | 0 |
| Crenarchaeota | 52 | 0 | 0 | 0 | 0 | 45 | 1 |
| Thaumarchaeota | 4 | 0 | 0 | 0 | 0 | 2 | 1 |

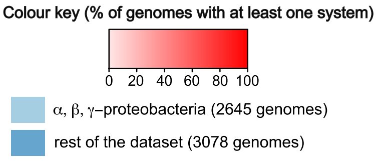


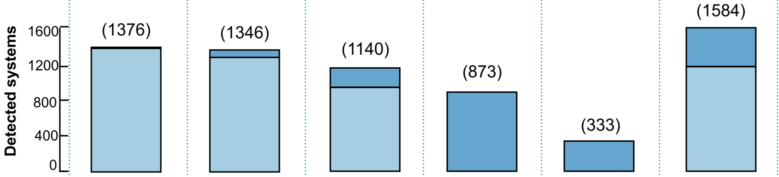
