## Supplemental Figure 15 for "Diversification of the type IV filament super-family into machines for adhesion, secretion, DNA transformation and motility"

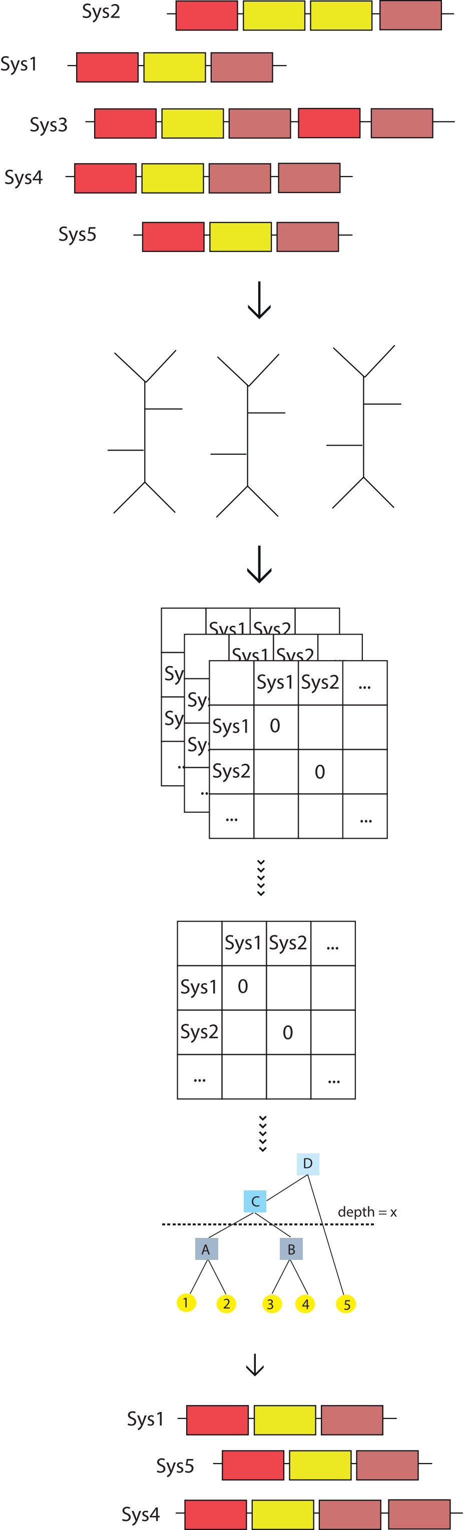


Step 1 : On the systems with at least ATPase and IM-platform protein

Step 2 : Inferring ML tree for each "core protein" family of the systems

Step 3a : Extracting patristic distances for all the trees

Step 3b : Inferring a bioNJ tree with the patristic distance

Step 3c : Cutting the tree at a fixed depth

Step 4 : Selecting one system for each cluster
