## Supplemental Figure 16 for "Diversification of the type IV filament super-family into machines for adhesion, secretion, DNA transformation and motility"

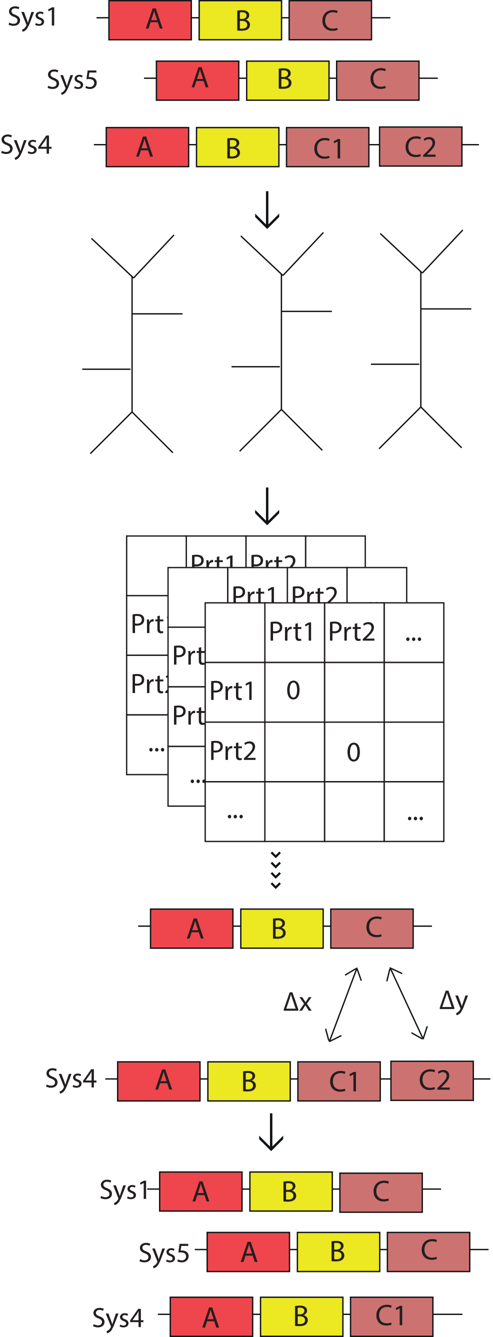


Step 1: Take all the systems representatives

Step 2 : Inferring ML tree for each "core protein" family of the systems

Step 3a : Extracting patristic distances for all the trees

Step 3b : Comparing the distances between the proteins in multi copies with the homolog in the closest systems

Step 4 : Choosing for each system the copy with the smallest distance
